# Average semivariance yields accurate estimates of the fraction of marker-associated genetic variance and heritability in complex trait analyses

**DOI:** 10.1101/2020.04.08.032672

**Authors:** Mitchell J. Feldmann, Hans-Peter Piepho, William C. Bridges, Steven J. Knapp

## Abstract

The development of genome-informed methods for identifying quantitative trait loci (QTL) and studying the genetic basis of quantitative variation in natural and experimental populations has been driven by advances in high-throughput genotyping. For many complex traits, the underlying genetic variation is caused by the segregation of one or more ‘large-effect’ loci, in addition to an unknown number of loci with effects below the threshold of statistical detection. The large-effect loci segregating in populations are often necessary but not sufficient for predicting quantitative phenotypes. They are, nevertheless, important enough to warrant deeper study and direct modelling in genomic prediction problems. We explored the accuracy of statistical methods for estimating the fraction of marker-associated genetic variance (*p*) and heritability 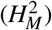 for large-effect loci underlying complex phenotypes. We found that commonly used statistical methods overestimate *p* and 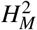. The source of the upward bias was traced to inequalities between the expected values of variance components in the numerators and denominators of these parameters. Algebraic solutions for bias-correcting estimates of *p* and 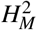 were found that only depend on the degrees of freedom and are constant for a given study design. We discovered that average semivariance methods, which have heretofore not been used in complex trait analyses, yielded unbiased estimates of *p* and 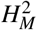, in addition to best linear unbiased predictors of the additive and dominance effects of the underlying loci. The cryptic bias problem described here is unrelated to selection bias, although both cause the overestimation of *p* and 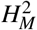. The solutions we described are predicted to more accurately describe the contributions of large-effect loci to the genetic variation underlying complex traits of medical, biological, and agricultural importance.

**Author summary:** The contributions of individual genes to the phenotypic variation observed for genetically complex traits has been an ongoing and important challenge in biology, medicine, and agriculture. While many genes have statistically undetectable effects, those with large effects often warrant in-depth study and can be important predictors of complex phenotypes such as disease risk in humans or disease resistance in domesticated plants and animals. The genes identified through associations with genetic markers in complex trait analyses typically account for a fraction of the heritable variation, a genetic parameter we called ‘marker heritability’. We discovered that textbook statistical methods systematically overestimate marker heritability and thus overestimate the contributions of specific genes to the phenotypic variation observed for complex traits in natural and experimental populations. We describe the source of the upward bias, validate our findings through computer simulation, describe methods for bias-correcting estimates of marker heritability, and illustrate their application through empirical examples. The statistical methods we describe supply investigators with more accurate estimates of the contributions of specific genes or networks of interacting genes to the heritable variation observed in complex trait studies.

## Introduction

The genetic variation observed in nature is frequently caused by genes with quantitative effects [1–7]. Their discovery and characterization has been a dominant feature of quantitative genetic studies in biology, evolution, agriculture, and medicine since the introduction of methods for genotyping DNA variants genome-wide [8–11], and the parallel development of statistical methods for finding associations between DNA variants and the underlying genes or quantitative trait loci (QTL) [2, 4, 5, 7, 12–16]. A significant breakthrough was achieved when Lander and Botstein [12] introduced ‘interval mapping’ and showed that genomes could be systematically searched to identify QTL in populations genotyped with a genome-wide framework of genetically mapped DNA markers. As genotyping technologies advanced and marker densities increased, genome-wide association study (GWAS) methods emerged to search genomes for genotype-to-phenotype associations by exploiting the historical recombination in populations [14, 15, 17–19]. The concept of genomic prediction emerged as a counterpart to GWAS, initially for estimating genomic-estimated breeding values (GEBVs) in domesticated plants and animals and later for estimating polygenic risk scores (PRSs) in humans and model organisms [20–23]. These technical advances precipitated a consequential shift in the study of quantitative traits from analyses of phenotypic variation limited and informed by pedigree or family data to genome-wide analyses of genotype-to-phenotype associations and genomic prediction informed by genotypic data [6, 7, 13, 16, 20, 24–31].

The phenotypic variation observed in a population is customarily partitioned into genetic and non-genetic components to estimate heritability, repeatability, and reliability of the quantitative traits under study [24, 25, 30, 32]. The genetic component can be caused by any number of genes with quantitative effects, even a single gene, but more often by multiple genes with a range of effects [31, 33–43]. For most quantitative traits, that number is unknown but presumed to be large and undiscoverable [3, 6, 7, 22, 32, 34]. Because genes with small effects are challenging to identify and validate, the ‘many genes with small effects’ hypothesis has been difficult to conclusively falsify [21, 22, 32]. Despite the uncertainty surrounding the identity, number, effects, and interactions of genes in the undiscovered fraction [6], three decades of complex trait analyses in humans, domesticated plants and animals, Drosophila, Arabidopsis, yeast, mice, zebrafish, and other organisms have shown that the ‘discovered’ genes are typically small in number, large in effect, and collectively only explain a fraction of the genetic variance 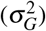 [13, 16, 28, 32, 34–36, 44, 45]. The unexplained fraction has been called ‘missing heritability’ [46–48].

The discovered genes in polygenic systems of genes are often necessary but not sufficient for predicting quantitative phenotypes, e.g., disease risk in humans or yield in domesticated plants and animals [3, 21, 34, 42, 44, 49]. There is a large body of evidence that the QTL effects for many quantitative traits are gamma family distributed, where the discovered genes are found in the upper or thin tail of the distribution above the threshold of statistical significance [34]. The presumption is that the lower or heavy tail of the gamma family distribution is caused by many genes with small effects, the chief tenet of the infinitesimal model of quantitative genetics [6, 26, 32, 50]. Genes with large effects often dominate the ‘non-missing heritability’, mask or obscure the effects of other quantitatively acting genes, and pleiotropically affect multiple quantitative phenotypes [16, 35, 39, 51], e.g., mutations in the *BRCA2* gene can have large effects, are incompletely penetrant, interact with other genes, and are necessary but not sufficient for predicting breast, ovarian, and other cancer risks in women [52]. The large-effect QTL BTA19 pleiotropically affects milk yield, protein yield, and productive life in Guernsey cattle (*Bos taurus*) [43], and branching and pigment genes (*BR, PHY*, and *HYP*) have large effects, interact, and pleiotropically affect several genetically correlated seed biomass traits in sunflower (*Helianthus annuus*) [53]. Despite decades of directional selection, loci with large effects often segregate (have not been fixed) in domesticated plant and animal populations [33, 34, 37, 38, 40, 54, 55]. The fractions of the genetic variances explained by *BRCA2*, BTA19, *BR, PHY*, and *HYP* were not reported in those studies. What fraction of the heritability for breast cancer risk, for example, can be explained by the known mutations in *BRCA2*? Our study explored the accuracy of methods for estimating that parameter.

Our surveys and others substantiate that the missing and non-missing fractions of the genetic variance are commonly either not estimated or inaccurately estimated in GWAS and other gene finding studies, e.g., the statistical significance of individual marker loci from sequential regression analyses are typically reported without correcting for the effects of other discovered marker loci through multilocus partial regression analyses or Type III ANOVA [17, 19, 22, 34, 56]. Such analyses are necessary for accurately assessing the statistical importance of the underlying gene and gene-gene interaction effects in a multilocus system, e.g., when multiple loci are identified by GWAS (sequential analyses of individual loci), their effects are more accurately estimated by simultaneous analysis using partial regression analysis approaches and even then can be upwardly biased [51]. The estimation problem we studied is intertwined with the broader problem of accurately describing multilocus systems of genes with large effects. We show that the discovered fraction of the genetic variance can be grossly overestimated and that the cause of the problem is a mathematical artifact in the expected values of variance components and their ratios. We revisited the problem of estimating the non-missing and missing fractions of heritability in candidate gene and other complex trait analyses, in part because of the systematic upward bias we discovered, in addition to inconsistencies in the methods commonly applied to the problem. The solutions to the problem presented here are straightforward and primarily applicable to the study of genes with large effects, especially those affecting the accuracy of genomic predictions for disease risk or breeding value [21, 43]. The optimum approaches for weighting or correcting for loci with large effects in genomic prediction are not completely clear; however, in artificial selection settings where the favorable alleles for discovered loci are unequivocally known, those alleles can be directly selected via marker-assisted selection (MAS) with genomic selection exerting pressure on unknown loci underlying the additive genetic variance not explained by the segregation of known large effect loci [54, 57–61].

Lande and Thompson [62] proposed the parameter 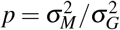 to estimate the discovered or non-missing fraction of the genetic variance, where 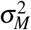 is the fraction of the genetic variance associated with statistically significant markers in linkage disequilibrium (LD) with genes or QTL affecting the trait under study (here QTL refers to a chromosome segment predicted to harbor a gene or genes affecting a quantitative trait). Similarly, marker heritability 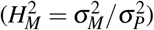 estimates the non-missing fraction of the phenotypic variance 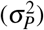 associated with statistically significant markers in LD with causal genes or QTL. Here a distinction needs to be made between 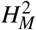 and genomic heritability, a parameter estimated by summing the effects of a dense genome-wide sample of markers, only some of which are predicted to be in LD with the underlying causal genes or QTL [27, 30, 63]. We are not proposing marker heritability as a replacement or substitute for genomic heritability but as a parameter for parsing out the non-missing fraction of heritability associated with discovered loci, especially loci like *BRCA2* and BTA19 [43, 52]. The genetic variance component 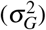 in these ratios can be estimated from pedigree or family information (as shown in our examples) or genomic information (as reviewed by [30] and [63]). For either, 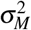 is simply the variance explained by marker loci with effects large enough to be statistically detected and important enough to be specifically studied and modeled, perhaps as fixed effects [22, 39, 40, 51, 61]. Despite a direct and logical connection to heritability, estimates of *p* and 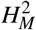 are seldom reported in complex trait studies, whereas genomic heritability estimates are commonly reported in genomic prediction studies [30, 34, 62].

Here we show that *p* and 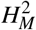 are often overestimated in complex trait analyses. The problem we discovered is unrelated to selection bias, the phenomena where the effects of discovered QTL are inflated by biased sampling from truncated distributions with small sample sizes [64–69], and unrelated to the upward biases known to arise in GWAS [70]. While selection bias is a well known and widely cited problem in complex trait analyses, we describe a previously unreported and cryptic source of bias in estimates of *p* and 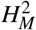. To identify the source of the bias and explore the problem in greater depth, we compared the accuracy of average marginal variance (AMV) [71, 72] and average semivariance (ASV) [73] methods for estimating *p* and 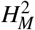. AMV is the acronym applied throughout this paper for the ANOVA and REML variance component estimation methods commonly described in textbooks and implemented in statistical software for the analysis of generalized linear mixed models (GLMMs), e.g., the ‘lme4’ R package and the SAS packages ‘GLM’ and ‘GLIMMIX’ [24, 25, 72, 74–79]. We introduced the average marginal variance terminology here to facilitate comparisons of the differences between AMV and ASV methods for estimating variance component *ratios*. The ASV methods we applied to the problem are extensions of those described by Piepho [73] for estimating the total variance and coefficient of determination (*R*^2^) in GLMM analyses. For the AMV and ASV analyses shown throughout this paper, REML was used to estimate the variance components [56, 72, 75, 79]. The source of the bias was discovered, however, through algebraic analyses of the expected mean squares (EMSs) from ANOVA. We describe that source and approaches for bias-correcting ANOVA or REML estimates of *p* and 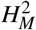 from the commonly applied AMV methods. We show that ASV methods directly yield unbiased estimates of *p* and 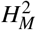 that are identical to bias-corrected AMV estimates. Finally, we discuss the connection of these random effects methods to the fixed effect methods commonly applied in QTL mapping and genome-wide association studies [51, 80, 81].

## Results and Discussion

### Overestimation of the Genetic Variance Explained by Markers in Linkage Disequilibrium With Causative Genes or QTL

The overestimation problem described here was originally discovered in a reanalysis of data from genetic studies in plants where REML estimates of 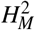 exceeded REML estimates of broad-sense heritability (*H*^2^) and REML estimates of *p* and 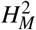 exceeded 1.0, the theoretical upper limit for these parameters (Table 1). We initially suspected that selection bias might be the culprit [68, 69, 82–84] but concluded that selection bias alone could not explain 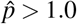 or 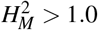. Although proof was lacking and the bias was non-obvious, we hypothesized that many estimates in the theoretical range (0.0 ≤ *p* ≤ 1.0) must also be upwardly biased. The proof was found through algebraic analyses of the ANOVA estimators of 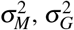, and 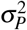 for balanced and unbalanced data (Appendices 2-4). Although variance components are commonly estimated using REML, as was done in the analyses shown throughout this paper, algebraic analyses of ANOVA expected mean squares (EMSs) identified the source of the bias and yielded explicit algebraic solutions for bias correcting ANOVA and REML estimates of *p* and 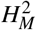.

**Table 1.**
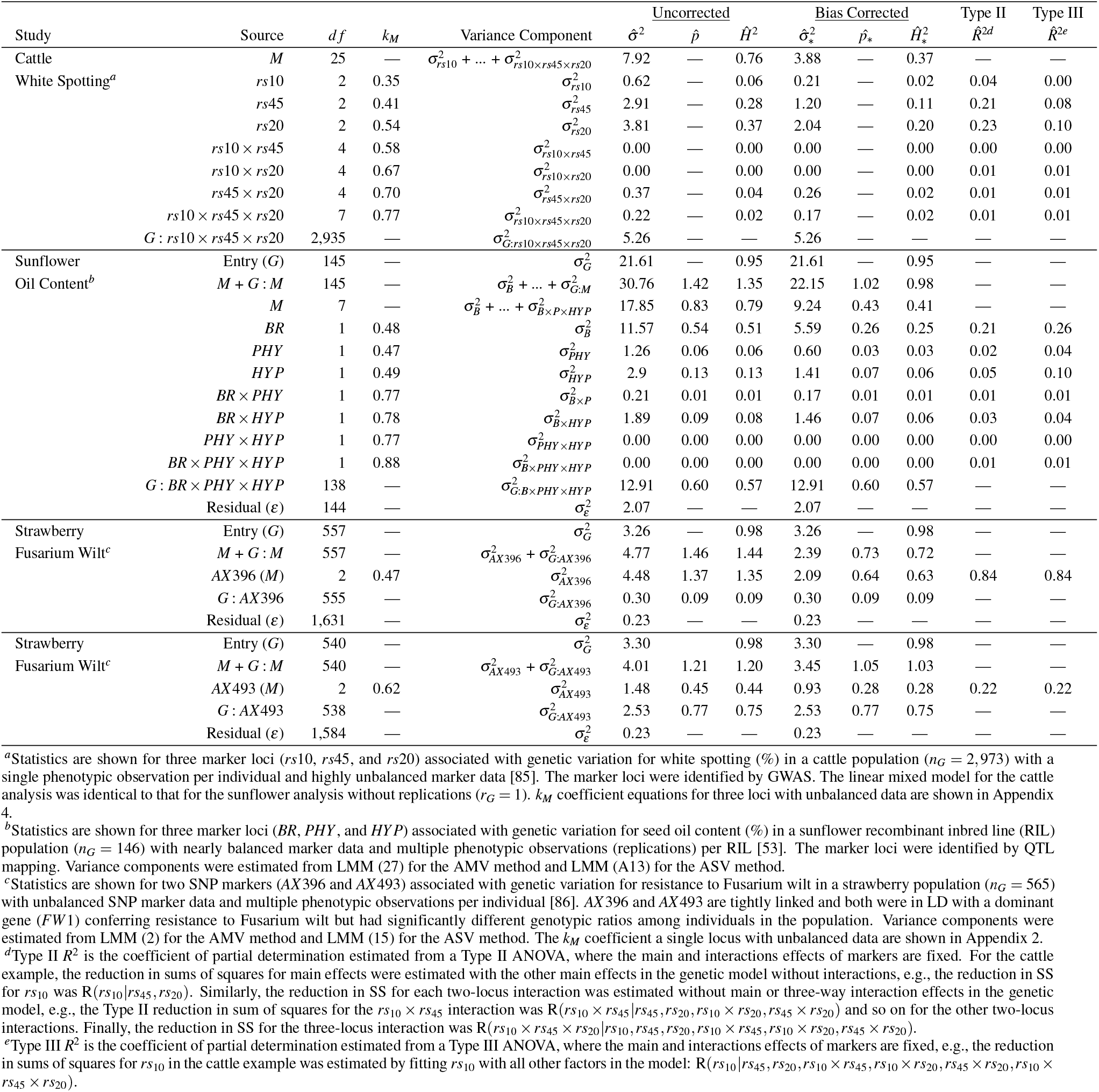
**REML estimates of marker-associated variance** 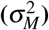, **the fraction of the genetic variance explained by markers** 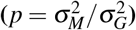, **and marker heritability** 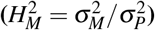 **from random marker effects analyses and coefficients of determination (***R*^2^**) from Type II and Type III fixed marker effects analyses for large effect loci identified in cattle, sunflower, and strawberry studies**.

The source of the bias was identified by expressing the estimator of as a function of the ANOVA estimators of 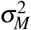 and 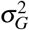 for balanced data and algebraically simplifying the equations. The linear mixed models (LMMs) and ANOVA estimators of the variance components needed to show this are described here. We start with the analysis of a single marker locus in an experiment where entries (e.g., individuals, families, or strains) are replicated, 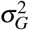 can be estimated, and the data for entries and markers are balanced. Extensions for one to three marker loci with unbalanced data are shown in Appendices 1-3. Two LMMs are needed for estimating 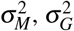, and 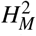. Consider a study where *n*_*G*_ entries are phenotyped for a normally distributed quantitative trait using a balanced completely randomized study design with *r*_*G*_ replications/entry, *n*_*M*_ marker genotypes/locus, and *r*_*M*_ replications/marker genotype. The LMM needed for estimating 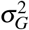 (the between entry variance component) is:

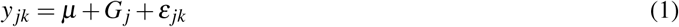

where *y* _*jk*_ is the *jk*^*th*^ phenotypic observation, *µ* is the population mean, *G*_*j*_ is the random effect of the *j*^*th*^ entry, *ε* _*jk*_ is the random effect of the *jk*^*th*^ residual, 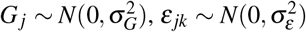, *j* = 1, 2, …, *n*_*G*_, and *k* = 1, 2, …, *r*_*G*_. Suppose entries are genotyped for a single marker locus (*M*) in linkage disequilibrium with a gene or QTL affecting the quantitative phenotype (*y* _*jk*_). The between entry source of variation from LMM (1) can be partitioned into marker (*M*) and entry nested in marker (*G* : *M*) sources of variation (this is the residual genetic variation among entries not explained by markers in the model). The LMM for estimating 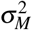 and 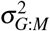 is:

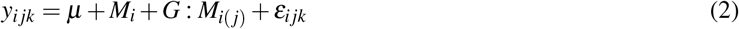

where *y*_*ijk*_ is the *i jk*^*th*^ phenotypic observation, *M*_*i*_ is the random effect of the *i*^*th*^ marker genotype at locus *M, G* : *M*_*i*(*j*)_ is the random effect of the *j*^*th*^ entry nested in the *i*^*th*^ marker genotype, *ε*_*ijk*_ is the random effect of the *i jk*^*th*^ residual, 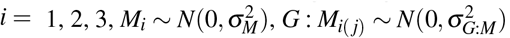, and 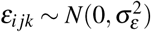.

The ANOVA estimator of the between-entry variance component 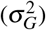 from LMM (1) with balanced data is:

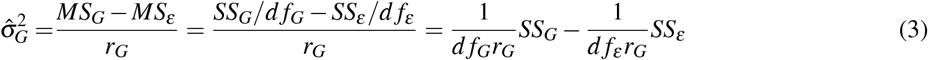

where *MS*_*G*_ = *SS*_*G*_*/df*_*G*_ is the between entry mean square, *SS*_*G*_ is the between entry sum of squares, *d f*_*G*_ = *n*_*G*_ − 1 is the between entry degrees of freedom, 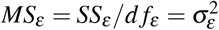 is the residual mean square, *SS*_*ε*_ is the residual sum of squares, *d f*_*ε*_ = *n*_*G*_(*r*_*G*_ − 1) −1 is the residual degrees of freedom, 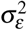 is the residual variance component, and *r*_*G*_ is the number of replications per entry [74]. The between-entry variance component has a theoretical genetic interpretation when entries are progeny with genetic relationships known from pedigrees, e.g., monozygotic twins, full-sib families, or recombinant inbred lines [24, 25, 30]. ANOVA estimators of the marker locus *M* and entry nested in *M* variance components from LMM (2) with balanced data are:

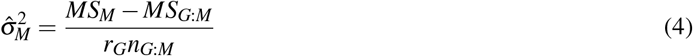

and

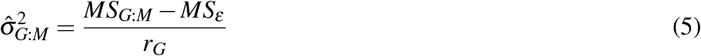

respectively, where *n*_*G*:*M*_ is the number of entries nested in each marker genotype, 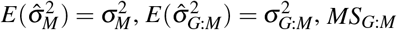, is the entry nested in *M* mean square, and *MS*_*M*_ is the mean square for marker locus *M*. The residuals in LMMs (1) and (2) are identical when the data are balanced 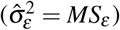. Hence, for a single marker locus with balanced data, the ANOVA estimator of *p* is:

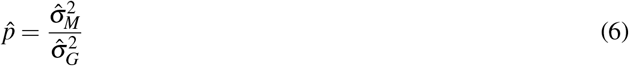

and the ANOVA estimator of broad-sense marker heritability on an entry-mean basis is:

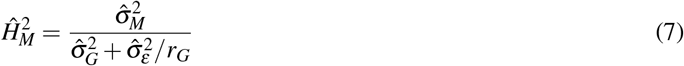

where 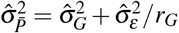 is the phenotypic variance on an entry-mean basis [25, 76].

The overestimation of *p* and 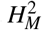 was not obvious from inspection of ANOVA estimators (6) and (7). The source of the bias was discovered by substituting *SS*_*M*_ + *SS*_*G*:*M*_ for *SS*_*G*_ in the ANOVA estimator of 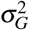 from (3) and simplifying:

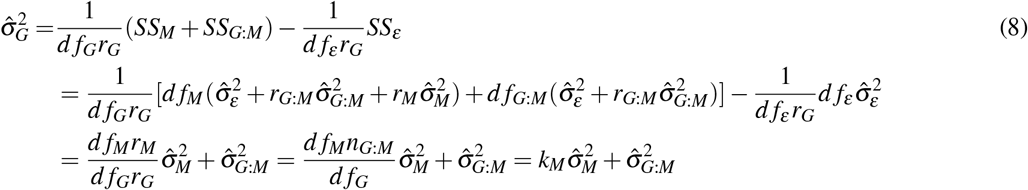

where the fraction *k*_*M*_ is source of the bias, 0 < *k*_*M*_ < 1, *r*_*M*_ is the number of replications per marker genotype, *n*_*G*:*M*_ is the number of entries nested in marker loci, *SS*_*M*_ is the marker sum of squares, *d f*_*M*_ is the marker degrees of freedom, *r*_*M*_ is the number of replicates of each marker genotype, *SS*_*G*:*M*_ is the entry nested in marker sum of squares, and *d f*_*G*:*M*_ is the entry nested in marker degrees of freedom. The term *k*_*M*_ in (8) depends on degrees of freedom and *n*_*G*:*M*_ and is hereafter referred to as the *k*_*M*_ bias coefficient, where the subscript *M* indexes the intralocus and interlocus effects of marker loci.

Equation (8) shows that the sum of ANOVA estimates of 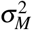 and 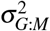 from LMM (1) are greater than the ANOVA estimate of 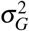 from LMM (2):

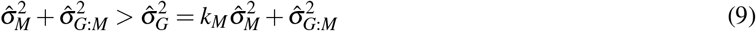

Although the SS for sources of variation in these LMMs are additive (*SS*_*M*_ + *SS*_*G*:*M*_ = *SS*_*G*_), the mean squares are not (*MS*_*M*_ + *MS*_*G*:*M*_ ≠ *MS*_*G*_). Because 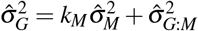, the sum 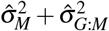 from LMM (2) overestimates 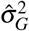 by a factor of 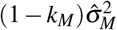. The ANOVA estimators of *p* and 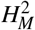 from analyses of LMMs (1) and (2) are upwardly biased because 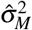 is multiplied by the fraction *k*_*M*_ in their denominators, and not the numerators:

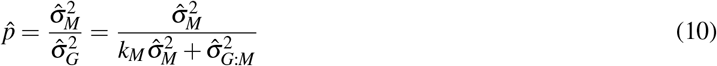

and

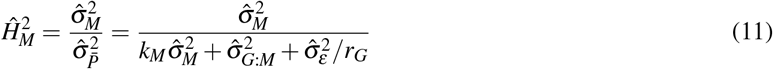

Substituting 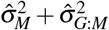 for 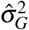 in the denominators of *p* and 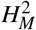 decreases but does not eliminate the bias because 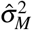 is multiplied by *k*_*M*_ in the denominator (Fig S1). For a single marker with balanced data, we found that:

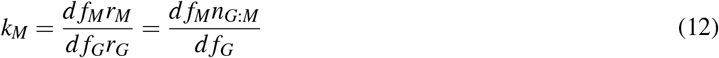

and

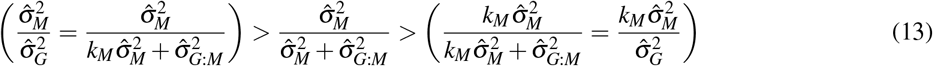

where 0 < *k*_*M*_ < 1. Hence, the bias is caused by the *k*_*M*_ multiplier in the expected values of the ANOVA estimators of *p* and 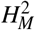. As shown later, simulation analyses confirmed that (9) and (13) accurately predict the upward bias caused by *k*_*M*_. Moreover, we concluded that the bias could be corrected by multiplying ANOVA or REML estimates of 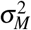 by *k*_*M*_ in the numerators of *p* and 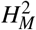 estimates.

### Genetic Models With Unbalanced Genotypic Data

We started with the special case of balanced data, which seldom arises in practice, but develop results here for the general case of unbalanced data. Following the same approach as that shown above for a single locus with balanced data, we found *k*_*M*_ coefficients for bias-correcting ANOVA and REML estimates of *p* and 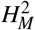 for analyses of one to three marker loci with unbalanced genotypic data (Appendices 1-3). For a single marker locus with unbalanced genotypic data, we found:

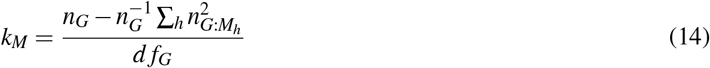

where *n*_*G*_ is the number of entries, *d f*_*G*_ are the degrees of freedom for entries, and 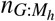is the number of entries nested in the *h*^*th*^ marker genotype (Appendix 1). This simplifies to (12) for a single marker locus with balanced genotypic data.

The *k*_*M*_ coefficients become slightly more complicated as the number of marker loci increases but nevertheless follow a predictable algebraic pattern, e.g., for a two locus genetic model, see equations (A10)-(A12) in Appendix 2. Similarly, for a three locus genetic model, see equations (A19)-(A25) in Appendix 3. *k*_*M*_ is greater (*k*_*M*_ bias is proportionally smaller) for interaction than main effects, e.g., for two marker loci, *k*_*M*1_ < *k*_*M*1×*M*2_ < 1 and *k*_*M*2_ < *k*_*M*1×*M*2_ < 1, where *k*_*M*1_ is the coefficient for *M*_1_, *k*_*M*2_ is the coefficient for *M*_2_, and 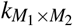 is the coefficient for the *M*_1_ × *M*_2_ epistatic interaction (Appendix 2). *k*_*M*_ for the two-locus interaction (*k*_*M*1×*M*2_) is larger than *k*_*M*_ for the individual marker loci (*k*_*M*1_ and *k*_*M*2_) because the denominator (*d f*_*G*_*r*_*G*_) is constant, whereas the numerators increase and approach the denominator as the degrees of freedom for marker effects increase. Therefore, the upward bias is proportionally smaller for the *M*1×*M*2 variance component than the *M*_1_ or *M*_2_ variance components for a two locus genetic model. Similarly, for a three locus genetic model, the upward bias is proportionally smaller for the *M*_1_ × *M*_2_ × *M*_3_ interaction variance component than the two-way interaction variance components (*M*_1_ × *M*_2_, *M*_1_ × *M*_3_, and *M*_2_ × *M*_3_). These results naturally extend to genetic models with more than three loci. Algebraic results are only shown for three marker loci because we found that the the *k*_*M*_ bias problem can be directly solved using average semivariance estimation methods when analyzing more complex genetic models (see below). Although certainly not limited to three marker loci, the methods described herein are primarily designed to study the effects of one to a few genes with large effects, e.g., *BRCA2* [52], BTA19 [43], and the examples shown in Table (1-2), and not to replace GWAS or QTL mapping.

### Study Designs Without Replications or Repeated Measures of Individuals or Families

LMMs (1) and (2) arise in study designs where entries, e.g., individuals, families, or strains, are replicated, e.g., in studies with domesticated plants, biological replicates of half-sib or full-sib families, doubled haploid or recombinant inbred lines, or testcross hybrids are commonly phenotyped [24, 25, 31, 76, 87, 88] (see the sunflower example in Table 1). These same LMMs apply to study designs for monozygotic twins in humans and other mammals and clonally replicated individuals in asexually propagated plants, e.g., cassava *(Manihot esculenta*), strawberry (*Fragaria* × *ananassa*), and apple (*Malus* × *domestica*) (see the strawberry examples in Table 1). The extension of the proposed *k*_*M*_ bias correction solutions to LMMs with repeated measures is straightforward and should have applications in studies where large effect loci are important determinants of the genetic variation underlying quantitative traits in both replicable and unreplicable organisms or populations [88–94].

When entries are unreplicated, the random error or residual source of variation in LMM (2) disappears (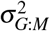 becomes the residual) and 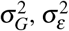, and *p* cannot be estimated; however, the marker heritability can be estimated using the phenotypic variance among unreplicated individuals 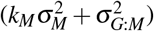. As before, this variance component ratio is upwardly biased by the factor *k*_*M*_ (see the cattle example in Table 1). Without the insights gained from the algebra shown in equations (10), (A3), (A9), and (A18), and Appendices 1-3, the bias would not be obvious unless one or more estimates of marker heritability exceeded 1, which only happens when the loci under study have very large effects. That was exactly how we originally discovered the bias problem in the first place (Table 1). The bias is systematic and ubiquitous but not immediately obvious when estimates fall within the expected range 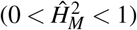. The same bias correction solutions we proposed for study designs with replications of entries can be applied in study designs where entries are unreplicated. When unreplicated entries are genotyped with a dense genome-wide of markers, 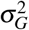 be estimated using a genomic or pedigree relationship matrix [92, 95–97], which yields an estimate of *p*.

### Average Semivariance Estimation Directly Solves the Bias Problem

The AMV methods proposed above for bias correcting ANOVA or REML estimates of *p* and 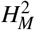 are straightforward to apply in practice because they are the methods widely described in textbooks and implemented in popular statistical software packages, e.g., the R package ‘lme4’ and SAS package ‘GLIMMIX’ [78, 98]. Here we show that the bias problem can be directly solved by applying average semivariance (ASV) estimation methods [73]. As before, we start by showing results for a single marker locus with balanced genotypic data. AMV notation and estimators are reformulated in matrix notation here to build the foundation for describing ASV notation and estimators. The input for both are the adjusted entry-level means 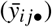 from LMM (1) stored in an *n*_*G*_-element vector. These are the best linear unbiased estimates (BLUEs) for entries [73, 99]. The LMM equivalent to (2) for the entry-level means analysis of the effect of a single marker locus (*M*) is:

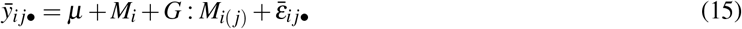

where 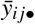 is the phenotypic mean for the *i j*^*th*^ entry, *µ* is the population mean, *M*_*i*_ is the random effect of the *i*^*th*^ marker genotype, 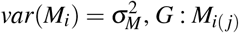 is the random effect of entries nested in *M*, 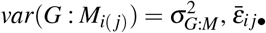 is the residual error, and 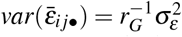. The residual variance-covariance matrix (*R*) is estimated in the first stage of a two-stage analysis [99–101]. The between-entry variance can be partitioned into 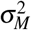 and 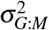 with individual variance-covariance matrices *G*_*c*_ defined by the genetic model, e.g., different main and interaction effects among marker loci.

The AMV estimator of the phenotypic (total) variance among observations for LMM (15) is:

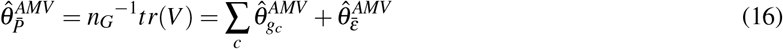

where *V* is the variance-covariance matrix of the phenotypic observations, *n*_*G*_ is the number of entries, *tr*(*V*) is the trace of *V*, 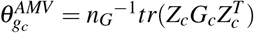 is the marginal variance explained by the *c*^*th*^ genetic factor in the model (e.g., *M* and *G* : *M*), *Z*_*c*_ are design matrices for the *c* genetic factors, 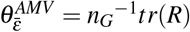 is the AMV estimator of the residual variance, and *R* is the residual variance-covariance matrix. The AMV estimator of the genetic variance among entries (*G*) is:

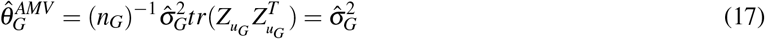

where 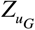 is a *n*_*G*_ identity matrix. From LMM (15), the AMV estimator of the variance associated with a single marker locus with balanced data is:

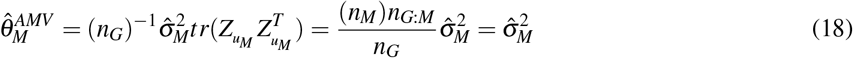

where 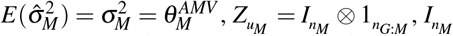 is a *n*_*M*_ identity matrix, 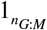 is an *n*_*G*:*M*_-element unit vector, and *u*_*M*_ is a vector of random effects for *M*. The AMV estimator of the variance associated with the residual genetic variation among entries nested in *M* is:

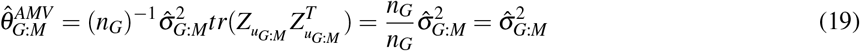

where *u*_*G*:*M*_ is a vector of random entry nested in *M* effects and 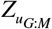 is a *n*_*G*_ identity matrix. Hence, the AMV estimators of 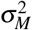 and 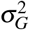 are identical to ANOVA estimators (4) and (5), respectively, with entry means as input for the former and original observations as input for the latter.

ASV, or the average variance of differences among observations, leads to a definition of the total variance that provides a natural way to account for the heterogeneity of variance and covariance among observations [73, 102]. ASV can be defined for any variance-covariance structure in a generalized LMM and allows for missing and unbalanced data [73]. The ASV estimator of total variance is half the average variance of pairwise differences among entries and can be partitioned into independent sources of variance, e.g., genetic and non-genetic or residual:

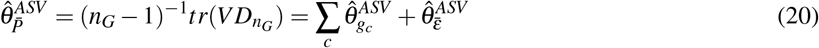

where 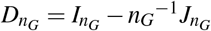 is the idempotent matrix used for column-wise mean-centering, 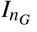 is an *n*_*G*_ × *n*_*G*_ identity matrix, and 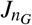 is an *n*_*G*_ × *n*_*G*_ unit matrix [73]. 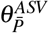 accounts for the variance and covariance of the phenotypic observations. From (20), 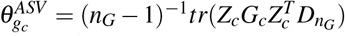 is the variance explained by the *c*^*th*^ genetic factor (*u*_*c*_), where *c* indexes genetic factors, the genetic factors are marker locus effects and entries nested in marker locus effects, and 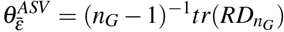 is the residual variance. The variance explained by the *c*^*th*^ genetic factor is 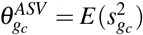, e.g., for a single marker locus *M*, 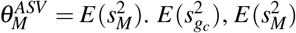, and the biases of these ASV estimators are defined in Appendix 4.

The ASV estimator of the genetic variance among entries (*G*) is:

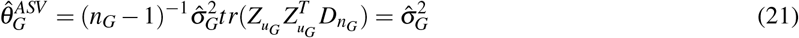

where 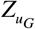 is a *n*_*G*_ identity matrix. Hence, from equations (8), (17), and (21), AMV and ASV estimators of the between-entry variance component 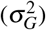 are equivalent 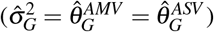. The ASV estimator of the variance associated with *M* is:

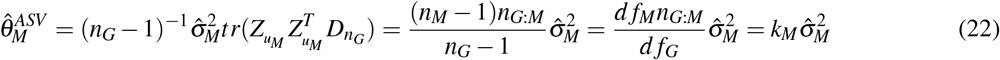

where *k*_*M*_ = *d f*_*M*_*n*_*G*:*M*_*/d f*_*G*_ is the bias correction coefficient, 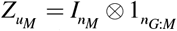, *df*_*G*_ = *n*_*G*_ −1, *df*_*M*_ = *n*_*M*_ −1, and *d f*_*G*:*M*_ = *d f*_*G*_ − *d f*_*M*_. This definition of the *k*_*M*_-bias coefficient is identical to the earlier definition with *r*_*G*_ factored out (see equation 12). Equation (22) shows that the ASV estimator of 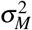 is corrected by the fraction *k*_*M*_, which correctly scales the estimate of 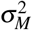 to the genetic variance and yields unbiased estimates of *p* and 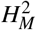. From equations (9) and (22), we found that 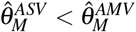 by the factor *k*_*M*_. The ASV estimator of the variance associated with *G* : *M* is:

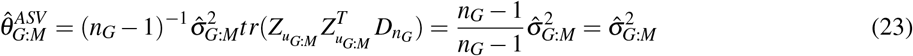

The ASV estimator of *p* for a single marker locus (*M*) is:

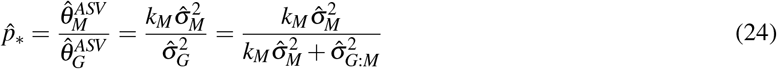

Similarly, the ASV estimator of 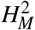 for a single marker locus is:

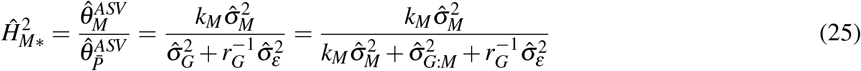

where 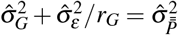 is the phenotypic variance on an entry-mean basis [25]. From these results, we found that:

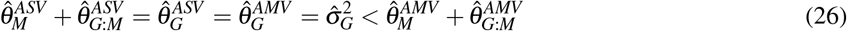

and showed that ASV estimators of *p* and 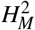 are unbiased (automatically corrected for *k*_*M*_).

### Computer Simulations Confirmed That ASV-REML Estimates of *p* and 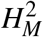 are Unbiased

Computer simulations confirmed that AMV-REML estimates of *p* (equation 6) and 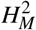 (equation 7) are upwardly biased by the factor *k*_*M*_ and that ASV-REML estimates of these parameters (equations 24) and 25) are unbiased (Fig 1 and 2). The mean of AMV-REML estimates of *p* and 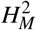 from 21 different simulation study designs (Table S1 were identical to those predicted by the *k*_*M*_ coefficients shown in Appendices 1-3. Several insights arose from the simulation analyses. First, the bias caused by *k*_*M*_ increased as 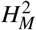 increased but was proportionally constant for different 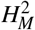 (Fig 1). These results show that the overestimation of *p* and 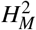 is greatest for genes and gene-gene interactions with large effects (Fig 1A-C). Their effects could be inflated by selection bias over and above *k*_*M*_ bias [67, 68, 82, 83]; hence, we concluded that *k*_*M*_-bias and selection bias could operate in combination to inflate estimates of the contribution of a locus to the heritable variation in a population (Appendix 2-4). Moreover, because the bias increases as the effect of the locus increases, we concluded that the overestimation problem is worst for large-effect QTL (Fig 1). Second, *k*_*M*_ bias was greater for unbalanced than balanced data (Fig 1D-E). The effect of unbalanced data was more extreme for the F_2_ simulation (Fig 1D) where the expected genotypic ratio was 1 *AA* : 2 *Aa* : 1 *aa* than for simulations where 10 or 33% of the observations were randomly missing for markers with roughly equal numbers of replicates/marker genotype (Fig 1E-F). Third, the F_2_ and missing data simulations further showed that the precision of estimates of these parameters decreased as the genotypic data imbalance increased. Even though bias-corrected AMV and ASV estimates of these parameters are unbiased, the sampling variances among the simulated F_2_ samples were larger than observed for the 10 and 33% missing data samples and yielded a small percentage of 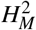 estimates slightly greater than 1.0 (Fig 1D). For the other simulation study designs (Fig 1A-C and E-F), none of the ASV estimates exceeded 1.0. The sample variances of *p* and 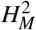 can be estimated using data resampling methods, e.g., bootstrapping [103], or the estimators we developed using the Delta method (Appendix 5). Equations (A44) and (A45) in Appendix 5 show that ASV estimates are more precise than AMV estimates by a factor of 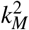. These predictions perfectly aligned with the empirical bootstrap estimates. Fourth, the relative biases were not affected by the number of replications of entries or the number of entries, although the precision of 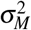 estimates increased as *n*_*G*_ and 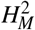 increased (Fig 2). Predictably, the number of entries(*n*_*G*_) dramatically affected the precision of estimates of 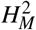 (Fig 2C-D). The relative biases were not affected by *r*_*G*_ or 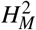; however, the sampling variances were strongly affected by 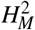 and decreased as 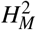 increased (Fig 2E-F and S2).

**Fig 1.**
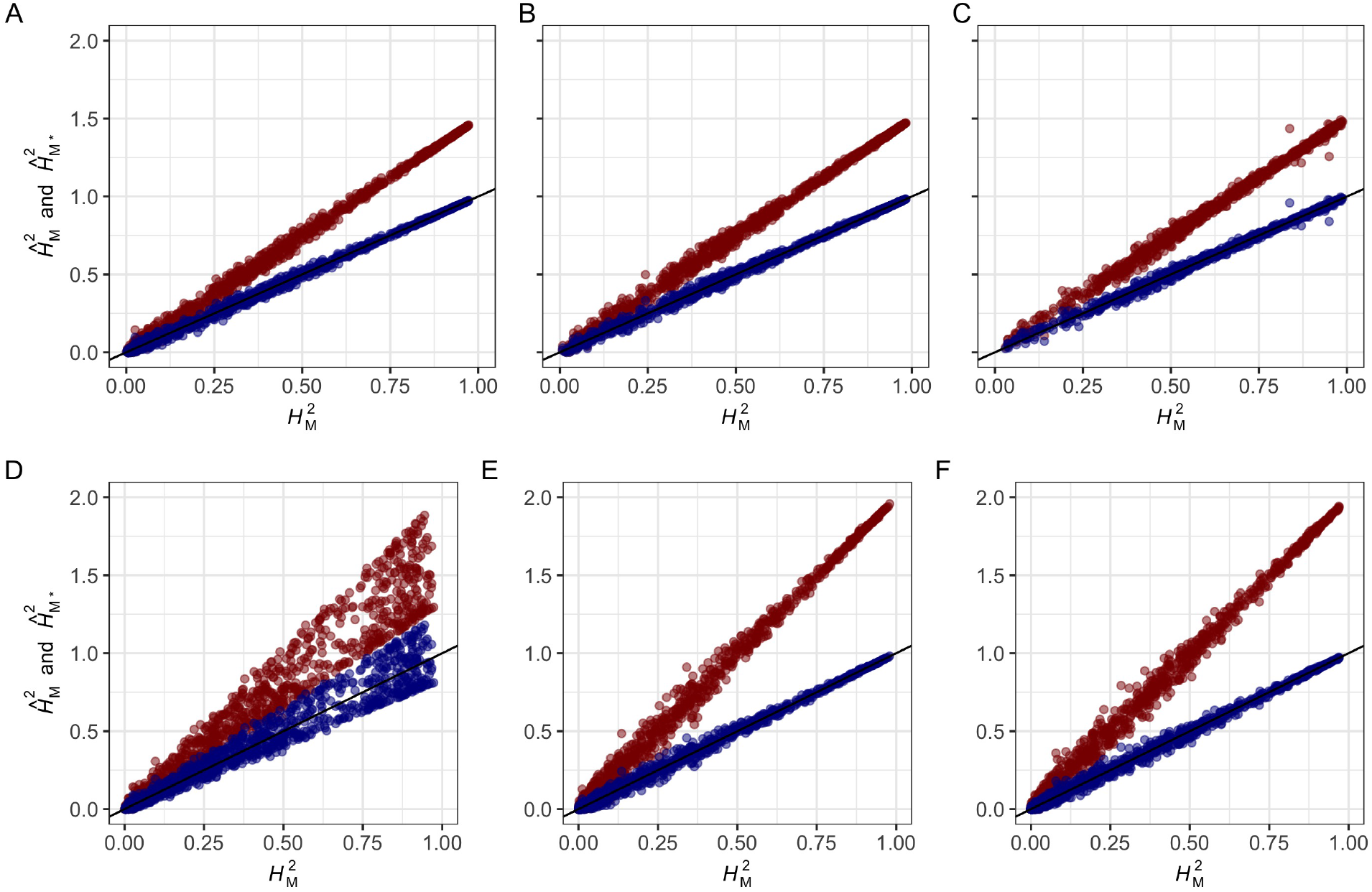
Accuracy of AMV and ASV Estimators of Marker Heritability. AMV and ASV estimates of 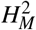 are shown for 1,000 segregating populations simulated for different numbers of entries (*n*_*G*_ individuals, families, or strains), five replications/entry (*r*_*G*_ = 5), true marker heritability 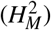 ranging from 0 to 1, and one to three marker loci with three genotypes/marker locus (*n*_*M*1_ = 3). AMV estimates of marker heritability (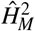; red highlighted observations) and ASV estimates of marker heritability (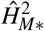; blue highlighted observations) are shown for: (A) one locus with balanced data for *n*_*G*_ = 540 entries (study design 1); (B) two marker loci with interaction (*M*1, *M*2, and *M*1 × *M*2) and balanced data for *n*_*G*_ = 540 (study design 2); (C) three marker loci with interactions (*M*1, *M*2, *M*3, *M*1 × *M*2, *M*1 × *M*3, *M*2 × *M*3, and *M*1 × *M*2 × *M*3) and balanced data for *n*_*G*_ = 540 (study design 3); (D) an population segregating 1:2:1 for one marker locus with *r*_*G*:*M*_ = 135 entries for both homozygotes and *r*_*G*:*M*_ = 270 heterozygous entries, and *n*_*G*_ = 540 (study design 4); (E) one locus with 10% randomly missing data among 540 entries (study design 5); and (F) one locus with 33% randomly missing data among 540 entries (study design 6). Study design details are shown in Table S1.

**Fig 2.**
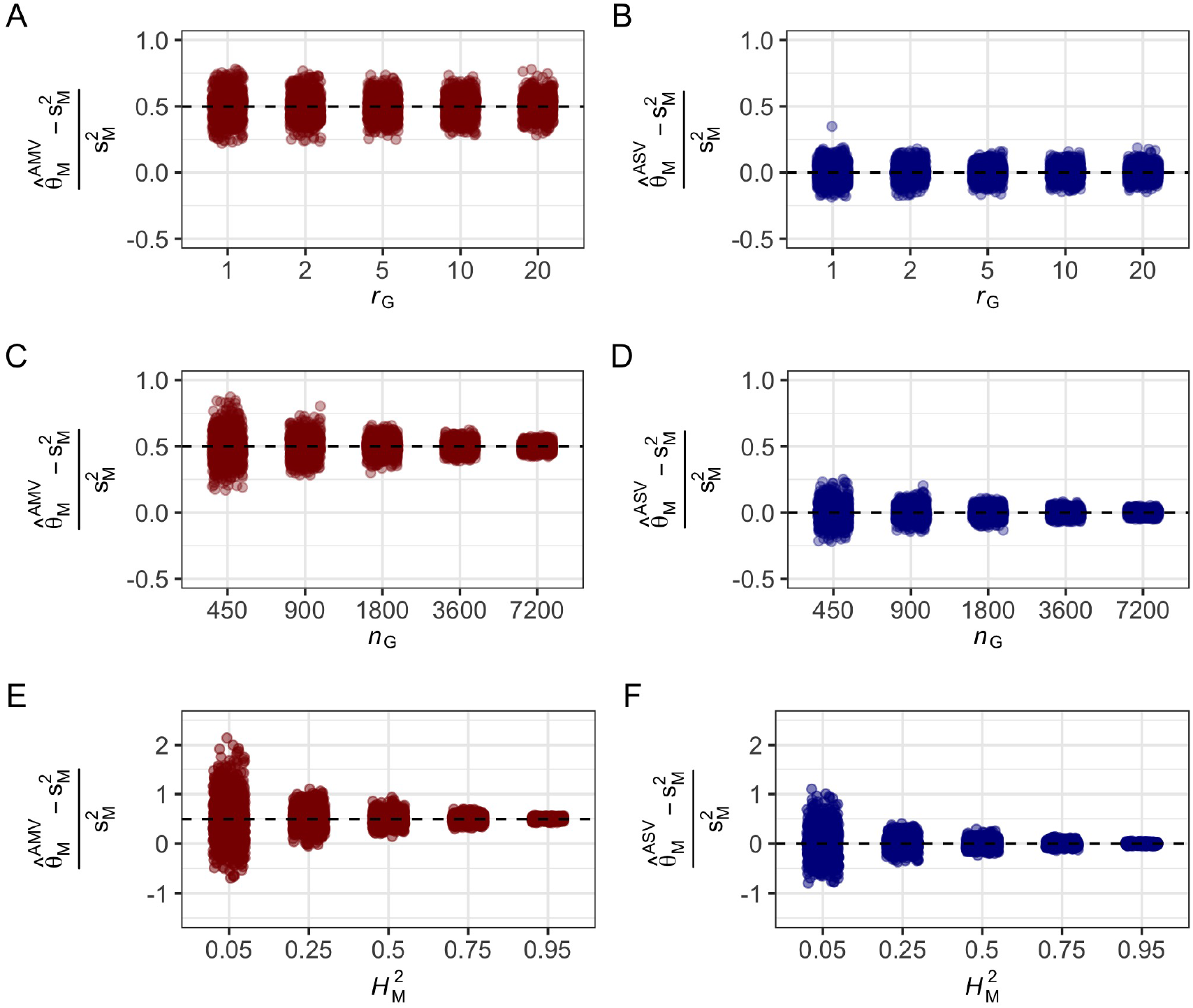
Effect of *r*_*G*_, *n*_*G*_, and 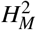 on the Relative Bias of AMV and ASV Estimators of 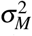. (A-B) Phenotypic observations were simulated for 1,000 populations segregating for a single marker locus with three genotypes (*n*_*M*_ = 3), *n*_*G*_ = 900 progeny, and *r*_*G*_ = 1, 2, 5, 10, or 20 (study designs 7-11). The marker locus was assumed to be in complete linkage disequilibrium with a single QTL that explains 50% of the phenotypic variance 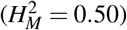. (A) Distribution of the relative biases of AMV estimates of 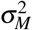 for different *r*_*G*_ . The relative bias 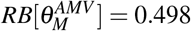 was identical for different *r*_*G*_. (B) Distribution of the relative biases of ASV estimates of 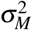 for different *r*_*G*_. The relative bias 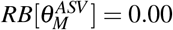 was identical for different *r*_*G*_. (C-D) Phenotypic observations were simulated for 1,000 populations segregating for a single marker locus with three genotypes (*n*_*M*_ = 3), five replications/entry (*r*_*G*_ = 5), and *n*_*G*_ = 450, 900, 1,800, 3,600, or 7,200 entries/population (study designs 12-16). The marker locus was assumed to be in complete linkage disequilibrium with a single QTL that explains 50% of the phenotypic variance 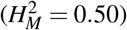. (C) Distribution of the relative biases of AMV estimates of 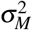 for different *n*_*G*_ . The relative bias 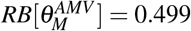 was identical across the variables tested. (D) Distribution of the relative biases of ASV estimates of 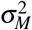 for different *n*_*G*_. The relative bias 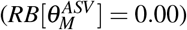 was identical across the variables tested. (E-F) Phenotypic observations were simulated for 1,000 populations segregating for a single marker locus with three genotypes (*n*_*M*_ = 3), five replications/entry (*r*_*G*_ = 5), and *n*_*G*_ = 450 entries/population. The marker locus was assumed to be in complete linkage disequilibrium with a single QTL that explains 5-95% of the phenotypic variance (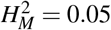 to 0.95 (study designs 17-21). (E) Distribution of the relative biases of AMV estimates of 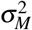 for different 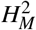. The relative bias 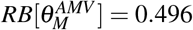 was identical across the variables tested. (F) Distribution of the relative biases of ASV estimates of 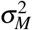 for different 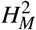. The relative bias 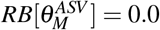 was identical across the variables tested.

### GWAS Example: A Single Marker Locus With Highly Unbalanced Genotypic Data

The bias-correction methods described above are illustrated here for highly unbalanced genotypic data from a GWAS experiment. Variance components were estimated for two SNP markers (*AX* 493 and *AX* 396) in LD with a gene (*FW* 1) conferring resistance to Fusarium wilt in a strawberry (*Fragaria* × *ananassa*) GWAS population (*n*_*G*_ = 564) genotyped with a genome-wide framework of SNP markers [86]. Both SNP markers had highly significant GWAS effects with − *log*_10_(*p*) = 6.61 × 10^−31^ for *AX* 493 and 2.95 × 10^−222^ for *AX* 396. Genotype frequencies were highly unbalanced for both markers with a scarcity of *AA* homozygotes (2.8%) for *AX* 396 (16AA: 177Aa: 371*aa*) and a 1 : 2 : 1 ratio for *AX* 493 (141AA: 282Aa: 141*aa*). For both loci, the minor allele frequency was *>* 0.05. The *k*_*M*_ for these data (*k*_*AX*493_ = 0.62 and *k*_*AX*396_ = 0.47) were calculated as shown in Appendix 1. The AMV-REML estimate of 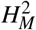 for *AX* 396 exceeded 1.0, a telltale sign of *k*_*M*_ -bias (Table 1). AMV-REML estimates of 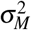 and 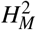 for both SNP markers were double or nearly double their bias-corrected ASV-REML estimates (Table 1). The bias-corrected estimate of marker heritability for *AX* 396 was 0.62, versus 1.33 for the uncorrected estimate. Even with bias-correction, the sum of ASV-REML estimates of 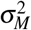 and 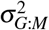 for *AX* 493 was slightly greater than the ASV-REML estimate of 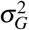. This result was consistent with finding for highly unbalanced marker genotypic data in our simulation studies where a certain fraction of bias-corrected estimates exceeded the theoretical limit for heritability because of decreased precision (Fig 1). The *k*_*M*_-bias problem would not necessarily have been detected in the analysis of *AX* 396 because the *p* and 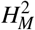 estimates fell within the expected range, e.g., 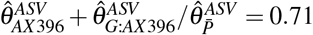 (Table 1). Although both SNP markers were closely associated with *FW* 1, they accounted for dramatically different fractions of genetic variance because of historic recombination and because neither are causal DNA variants or in complete LD with causal DNA variants [17, 19, 86, 104].

### QTL Mapping Example: Three Marker Loci With Slightly Unbalanced Genotypic Data

Statistics are shown here for an analysis of three marker loci (*BR, PHY*, and *HYP*) affecting seed oil content in a sunflower (*Helianthus annuus*) RIL population using LMM (27) [53]. The genotypic data were only slightly unbalanced and the three marker loci were identified by QTL mapping. The *k*_*M*_ needed for bias-correcting AMV-REML estimates of *p* and 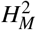 are shown in Appendix 3 (Table 1). The AMV-REML estimates of *p* and 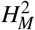 were nearly double the bias-corrected ASV-REML estimates, e.g., the AMV-REML estimate of 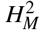 for the three-locus genetic model (0.79) was nearly two-fold greater than the ASV-REML estimate (0.41) (Table 1). Similarly, the AMV-REML estimate of *p* for the *BR* locus (0.54) was slightly more than double the bias-corrected (ASV-REML) estimate (0.26). Hence, the uncorrected REML estimates of *p* and 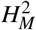 grossly inflated the predicted contributions of the three marker loci to genetic variation for seed oil content (Table 1).

### GWAS Example: Three Marker Loci With Unbalanced Genotypic Data and Unreplicated Entries

The application of bias-correction is illustrated here for a genetic model with three marker loci, highly unbalanced genotypic data, and a single phenotypic observation per individual— 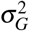 and *p* could not be estimated for this example because individuals were unreplicated. Variance components were estimated for three SNP markers (*rs*10, *rs*45, and *rs*20) on chromosomes 2, 6, and 22, respectively, affecting white spotting (%) in a Holstein–Friesian cattle (*Bos taurus*) population (*n*_*G*_ = 2, 973) [85]. These SNP markers had the largest effects among those predicted to be in LD with genes affecting white spotting. The genotypic frequencies were 50AA: 586Aa: 2, 337*aa* for *rs*10, 78AA: 736Aa: 2, 159*aa* for *rs*45, and 237AA: 976Aa: 1, 760*aa* for *rs*20. The *k*_*M*_ for these data (*k*_*rs*10_ = 0.35, *k*_*rs*45_ = 0.41, and *k*_20_ = 0.54) were calculated as shown in Appendix 3. The uncorrected AMV-REML estimate of 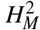 for the three-locus genetic model (0.76) was substantially greater than the bias-corrected ASV-REML estimate (0.37) (Table 1). Similar differences were observed for the three marker loci.

### Candidate Gene Analysis: Fixed or Random, BLUE or BLUP?

Our study was partly motivated by inconsistencies in the statistical approaches applied in candidate gene and other complex trait analyses when testing hypotheses and fitting genetic models for multiple large-effect loci. With the high densities of genome-wide markers commonly assayed in gene finding studies, investigators often identify markers tightly linked to candidate or known causal genes, as exemplified by diverse real world examples [17, 19, 33, 34, 37, 38, 40, 42, 43, 52, 54, 105]. The candidate marker loci are nearly always initially identified by genome-wide searches using sequential (marker-by-marker) approaches [56, 72, 75, 79, 106, 107]. Complicated and often misunderstood problems arise in the estimation and interpretation of statistics from sequential fixed effect analyses when the data are unbalanced [79, 108, 109]. Most importantly, there are multiple model fitting and analysis options (Type I, II, and III ANOVA) and the reduction in error sums of squares (SSE), test statistics, and parameter estimates differ among them, a problem that disappears when the data are balanced or when single large effect loci are discovered [79, 108–110]. Our review of the literature uncovered substantial variation and inconsistencies in the statistical approaches applied to the problem of fitting multilocus genetic models, testing multilocus genetic hypotheses, and estimating best linear unbiased estimates (BLUEs), e.g., estimated marginal means from a fixed effects analysis of marker loci.

The problems that arise in fixed effect analyses of unbalanced data profoundly affect parameter estimates and statistical inferences but have not been universally recognized or addressed in complex trait analyses [79, 109]. We reanalyzed the cattle and sunflower examples with markers as fixed effects (Table 2) to show this, illustrate the challenges and nuances of fixed effects analyses of unbalanced data, and facilitate comparisons between random and fixed effects analyses of marker loci [56, 56, 75, 79, 106–109]. Following the discovery of statistically significant marker-trait associations from a marker-by-marker genome-wide scan, the natural progression would be to analyze multilocus genetic models where the effects of the discovered loci are simultaneously corrected for the effects of other discovered loci [79, 109], as shown in our multilocus analysis examples (Table 1-2). This is straightforward when the genotypic data are balanced or nearly balanced (as in the sunflower example) but more complicated and convoluted when the genotypic data are unbalanced (as in the cattle example) [75, 79, 108, 109]. Although methods for fixed effect analyses of factorial treatment designs (multilocus genetic models) with unbalanced data are well known [56, 79, 106, 107, 109], there are several model fitting and parameter estimation variations that can lead to dramatically different parameter estimates and statistical inferences. This is perfectly illustrated by the cattle example where the coefficients of determination (analogous but not identical to 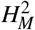) from Type I, II, and III analyses were substantially different from each other and from 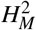 estimates from the random effects analysis (Table 1-2). The differences and ambiguities among the different fixed effects approaches disappear when the random effects approach is applied to the problem.

**Table 2.**
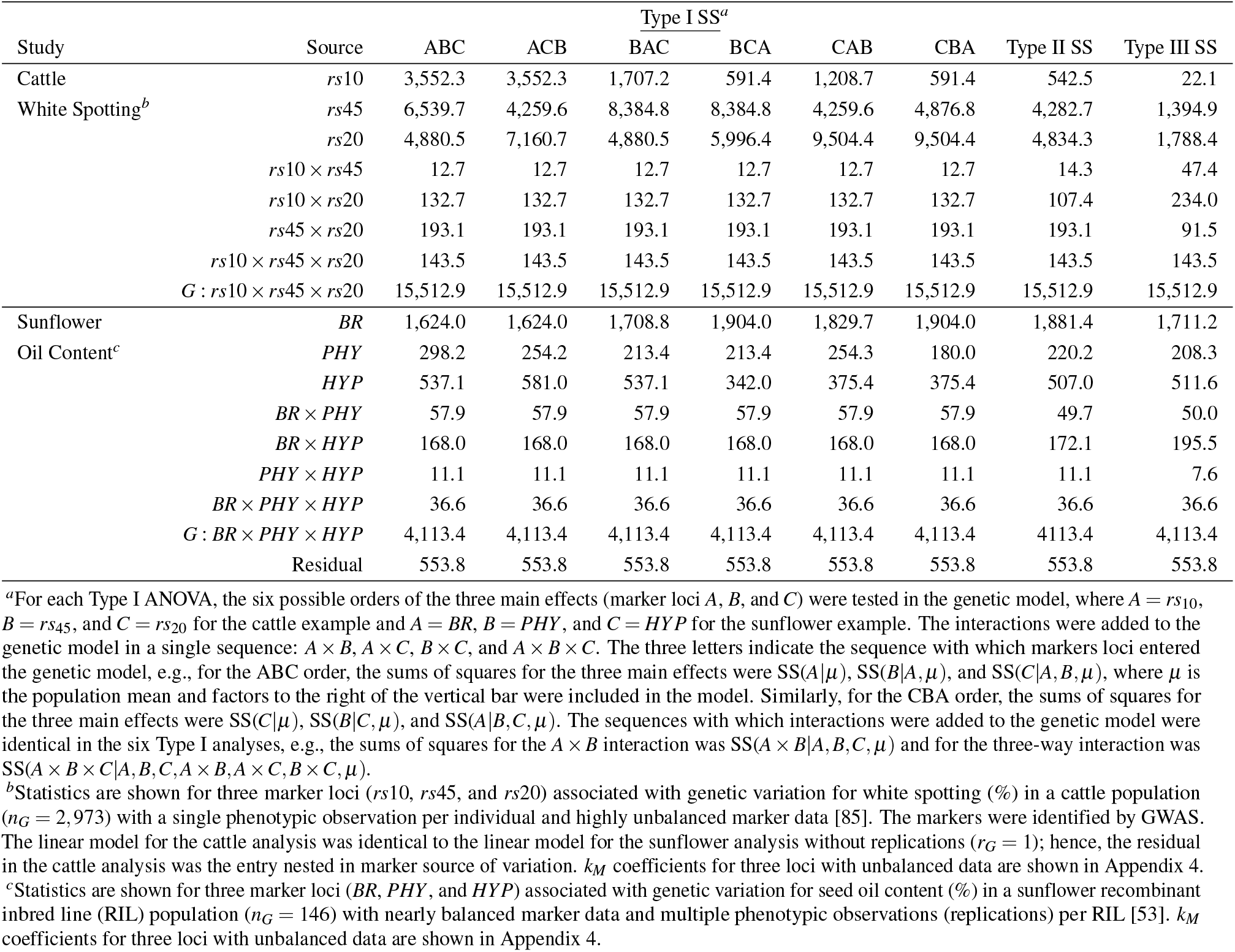
**Type I, II, and III sums of squares for fixed effect analyses of markers associated with QTL identified in GWAS and QTL mapping experiments in cattle and sunflower.**

The analysis of markers as random effects in a multilocus analyses of known or candidate genes with large effects with ASV, although historically uncommon, simultaneously yields unbiased estimates of the variance component ratios investigated in the present study (*p* and 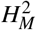) and best linear unbiased predictors (BLUPs) of the additive and dominance effects of the causative loci identified by marker associations, in addition to solving the often ambiguous problems that arise in fixed effects analyses of unbalanced data [32, 75, 77, 79, 109, 110]. As discussed in depth below and illustrated through a reanalysis of the cattle and sunflower examples (Table 2), the random effects approach we described (ASV with REML estimation of the variance components) yields accurate estimates of the underlying genetic parameters (variance component ratios and BLUPs of marker effects) from a *single* unambiguous generalized linear mixed model analysis, whereas wildly different parameter estimates can arise among the multitude of fixed effects analyses that investigators might elect to apply in practice when the underlying genotypic and phenotypic data are unbalanced (Tables 1-2).

As substantiated by our simulation analyses (Fig 1-2), ASV with REML estimation of the underlying variance components yields accurate estimates of *p* and 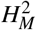 for marker loci and interactions between marker loci, both individually and collectively, and BLUPs of the the additive and dominance effects of marker loci [76, 110–112]. When the genotypic data are unbalanced, the order with which marker and marker × marker effects enter the genetic model profoundly affects parameter estimates and statistical inferences in fixed effect analyses [56, 72, 74, 113]. To illustrate this, the main effects of marker loci A, B, and C were estimated for the six possible Type I ANOVA orders of the three loci (ABC, ACB, BAC, BCA, CAB, and CBA) (Table 2). Predictably, the reduction in the error sums of squares for a particular locus differed for each Type I order in the cattle example: the Type I SS ranged from 591.4 to 3,552.3 for *rs*10, 4,880.5 to 9,504.4 for *rs*20, and 4,259.6 to 8,384.8 for *rs*45. The *R*^2^, or PVE, estimates for marker loci were radically different among the six Type I ANOVA and Type II and III analyses. The Type I SS were, in addition, significantly greater than the Type III SS for nearly every factor. Although Type III statistics are commonly estimated and reported in analyses of factorial treatment designs with unbalanced data, there are compelling arguments for estimating Type II statistics [106, 107]; nevertheless, as we have argued, the fixed effects approach is unnecessary.

Broadly speaking, the large effect loci segregating in a population are typically necessary but not sufficient for predicting genetic merit or disease risks but are often important enough to warrant deeper study and, in animal and plant breeding, direct selection via MAS or direct modelling in genome selection applications [21, 32, 57]. The BLUP (random marker effects) approach we applied was designed to align the study of loci with large and highly predictive effects with the BLUP approaches commonly applied to genomic prediction problems that are agnostic or indifferent to the effects of individual loci, the so-called “black box” of genomic prediction [6, 7, 20, 21, 88, 114–118]. The predictive markers associated with large effect marker loci can be integrated into the genome-wide framework of marker loci applied in genomic prediction or incorporated as fixed effects when estimating GEBVs or PRSs [21, 54, 57–61]. One of the greatest strengths of the random effects (BLUP) approach is that the genetic parameters can be estimated from a single REML analysis free of the challenges and uncertainty associated with the fixed effects model building process [79, 106, 107, 109]. Finally, if our conclusions are correct, the complex trait analysis literature is riddled with overestimates of the genotypic and phenotypic variances explained by specific genes or QTL.

## Materials and Methods

### Simulation Studies

We used computer simulation to estimate the bias and assess the accuracy of uncorrected and bias-corrected REML estimates of *p* and 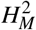 for 21 study designs (Table S1; Appendix 4). Phenotypic observations (*y*_*i jk*_) for LMMs (1) and (2) were simulated for *n*_*M*_ = 3 genotypes/marker locus and 21 combinations of study design variables (*n*_*G*_, *r*_*G*_, *r*_*M*_, and *H*^2^) with balanced or unbalanced data (Table S1). Simulations were performed to assess the accuracy of REML estimates of *p* and 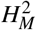 for 21 study designs with 1,000 replicates per study design (Table S1). The phenotypic observations for each sample were obtained by generating random normal variables for entries, markers, and residuals using the R function *rnorm()* with known means and variances [119] as described by [120, 121]. The simulated random effects of entries, markers, and replications in LMMs (1) and (2) were summed to obtain *n* = *n*_*G*_*r*_*G*_ phenotypic observations for each study design. Variance components for the random effects in LMMs (1) and (2) were estimated using the REML function implemented in and assess the accuracy of AMV and ASV estimators of *p* and 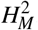. For study designs 1-6, the true marker heritability randomly varied from 0 to 1. Study designs 1-6 demonstrate how different numbers of marker loci (*m*) and unbalanced data affect estimates of *p* and 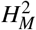 (Fig 1; Table S1). For study designs 5 and 6, we randomly deleted 10 and 33% of the phenotypic observations, respectively, to create unbalanced data. For study designs 7-21, the true variances of the independent variables were fixed for all samples, which allowed us to estimate the bias and relative bias associated with the different estimators (the biases are shown in Appendix 1). Study designs 7-21 illustrate how *r*_*G*_, *n*_*G*_, and 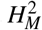 affected the biases and relative biases of *p* and 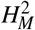 (Fig 2; Table S1; Appendix 4). The variance components were estimated using REML in the *lme4::lmer()* v1.1-21 [78] package in R v4.0.2 [119]. We estimated the sample variances of AMV and ASV estimates of *p* for each study design (Table S1). Finally, we developed estimators of the sampling variances of *p* and 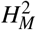 using the delta method [25, 122], as shown in Appendix 5.

### Estimation Examples

To illustrate the application of bias-correction methods and the differences between AMV and bias-corrected AMV estimates of *p* and 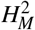, we reanalyzed data from a GWAS study in cattle (*Bos taurus*), a QTL mapping study in oilseed sunflower (*Helianthus annuus* L.) [53], and a GWAS study of Fusarium wilt resistance in strawberry (*Fragaria* × *ananassa* Duchesne ex Rozier) [86]. For the sunflower study, two replications (*r*_*G*_ = 2) of *n*_*G*_ = 146 recombinant inbred lines (RILs) were phenotyped for seed oil concentration (*g/kg*) and genotyped for three marker loci (*BR, PHY*, and *HYP*) with two homozygous marker genotypes/locus [53]. For the cattle study, unreplicated entries (*r*_*G*_ = 1; *n*_*G*_ = 2, 973) were phenotyped for white spotting (%) and genotyped for three marker loci (*rs*10, *rs*45, *rs*20) with three marker genotypes per locus [85]. LMM (2) expanded to three marker loci with all possible interactions among marker loci is:

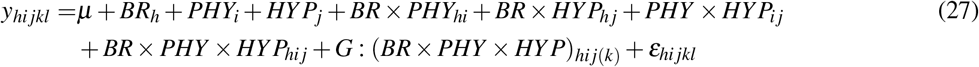

where *BR*_*h*_ is the *h*^*th*^ effect of the *BR* locus, *PHY*_*i*_ is the *i*^*th*^ effect of the *PHY* locus, *HYP*_*j*_ is the *j*^*th*^ effect of the *HYP* locus, *G* : (*BR* × *PHY* × *HYP*)_*hi j*(*k*)_ is the *k*^*th*^ effect of entries nested in the *hi j*^*th*^ *BR* × *PHY* × *HYP* interaction, and *ε*_*hi jkl*_ is the *hi jkl*^*th*^ residual effect. The data for RILs were balanced, whereas the data for marker genotypes were slightly unbalanced. Each of the eight *BR* × *PHY* × *HYP* homozygotes were observed in the RIL population; however, the number of entries nested in each marker genotype (*n*_*G*:*M*_) varied from *n*_*G*:*BR*_ = 81 : 65, *n*_*G*:*PHY*_ = 60 : 86, and *n*_*G*:*HYP*_ = 70 : 76. Variance components for LMMs (1) and (27) were estimated using the REML method in *lme4::lmer()* [78]. The marker-associated genetic variances for individual marker loci and two- and three-way interactions among marker loci were bias-corrected using the formula described in Appendix 4.

For the strawberry study, four replications *(r*_*G*_ = 4) of 565 entries (*n*_*G*_ = 565) from a genome-wide association study (GWAS) were phenotyped for resistance to Fusarium wilt and genotyped for single nucleotide polymorphism (SNP) markers in LD with *FW* 1, a dominant gene conferring resistance to *Fusarium oxysporum* f.sp. *fragariae*, the causal pathogen [86]. The replications were asexually propagated clones of individuals; hence, the expected causal variance among individuals was equal to the total genetic variation in the population, analogous to monozygotic twins [25]. Genetic parameters were estimated for two SNP markers (*AX* 493 and *AX* 396) that were tightly linked to *FW* 1 [86]. The genotypic data for both markers were highly unbalanced. Genotype numbers were 141 *AA* : 282 *Aa* : 141 *aa* for *AX* 493 and 16 *AA* : 177 *Aa* : 371 *aa* for *AX* 396, where *A* and *a* are alternate SNP alleles. The variance components were estimated for LMMs (1) and (2) using REML method implemented in the R package *lme4::lmer()* [78]. REML estimates of the marker-associated genetic variances for both marker loci were bias-corrected using the approach described in Appendix 2.

For the cattle study, we used a model similar to (27) for the analysis. However, because entries are unreplicated in this experiment, we cannot include the entries nested in the three-way marker interaction *(G* : *M*) term because it has the same levels as the residual. The LMM for this case study is:

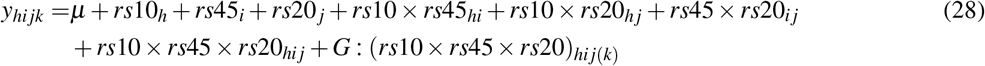

where *rs*10_*h*_ is the *h*^*th*^ effect of the peak SNP (*rs*109979909) on chromosome 2, *rs*45_*i*_ is the *i*^*th*^ effect of the peak SNP on chromosome 6 (*rs*451683615), *rs*20 _*j*_ is the *j*^*th*^ effect of the peak SNP on chromosome 22 (*rs*209784468), and *G* : (*rs*10 × *rs*45 × *rs*20)_*hi j*(*k*)_ is the *hi j*(*k*)^*th*^ residual effect comprising residual genetic effects *G* : *M* and residual error. In this experiment, there were *k* entries and *k* observations, and because of this we cannot fit LMM (1) without incorporating pedigree or genomic relatedness. In this single case, we estimate 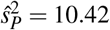 from the log transformed data to use in the denominator of 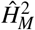.

We used the SAS package PROC GLM [77] for Type I and III analyses of the sunflower and cattle data with marker loci as fixed effects. Type I analyses were done for the six possible orders of main effects (ABC, ACB, BAC, BCA, CAB, and CBA) and a single order for marker × marker interactions (A × B, A × C, B × C, and A × B × C), where A, B, and C are the three marker loci (factors). For the ABC order, the reduction SS for the main effects were R(A | *µ*), R(B |*µ*, A), and R(C | *µ*, A, B), where *µ* is the population mean. Similarly, for the ACB order, the reduction SS for the main effects were R(A | *µ*), R(C |*µ*, A), and R(B | *µ*, A, C), and so on for the other four orders (BAC, BCA, CAB, and CBA). C, A × B, A × C, B × C, A × B × C), the reduction SS for the main effect of B was R(B | B, C, A × B, A × C, B × C, A × B × C). For comparison, the Type III reduction SS for the main effects were R(A | *µ*, B, C, A × B, A × C, B × C, and A × B × C), R(B | *µ*, A, C, A × B, A × C, B × C, and A × B × C), and R(C | *µ*, A, B, A × B, A × C, B × C, and A × B × C).

### Data Availability

The custom R scripts for reproducing our simulations have been deposited in a public GitHub repository [123]. The simulated data shown in Figs 1 and 2 have been deposited in a public Zenodo repository [124].

## Abbreviations

AMV: average marginal variance
ANOVA: analysis of variance
ASV: average semivariance
GEBV: genome estimated breeding value
GWAS: genome-wide association study
LD: linkage disequilibrium
LSR: least squares regression
MAS: marker-assisted selection
QTL: quantitative trait locus
REML: restricted maximum likelihood
SNP: single nucleotide polymorphism
WGR: whole-genome regression

## Acknowledgements

The authors thank the anonymous reviewers for comments and suggestions that improved the manuscript.

## Author Contributions

**Conceptualization:** Steven J. Knapp, William C. Bridges, Mitchell J. Feldmann, Hans-Peter Piepho

**Data curation:** Mitchell J. Feldmann

**Formal Analysis:** Mitchell J. Feldmann, Hans-Peter Piepho, Steven J. Knapp, William C. Bridges

**Funding Acquisition:** Steven J. Knapp and Hans-Peter Piepho

**Investigation:** Mitchell J. Feldmann, Hans-Peter Piepho, Steven J. Knapp

**Methodology:** Mitchell J. Feldmann, Hans-Peter Piepho, Steven J. Knapp, William C. Bridges

**Project administration:** Steven J. Knapp

**Resources:** Steven J. Knapp and Mitchell J. Feldmann

**Software:** Mitchell J. Feldmann

**Supervision:** Steven J. Knapp and Mitchell J. Feldmann

**Validation:** William C. Bridges, Hans-Peter Piepho, Mitchell J. Feldmann, Steven J. Knapp

**Visualization:** Mitchell J. Feldmann, Steven J. Knapp, Hans-Peter Piepho

**Writing – original draft preparation:** Mitchell J. Feldmann, Steven J. Knapp, Hans-Peter Piepho

**Writing – review & editing:** Mitchell J. Feldmann, Hans-Peter Piepho, Steven J. Knapp, William C. Bridges

## Appendices

### A1 ASV Estimator of the Fraction of the Genetic Variance Associated With a Single Marker Locus for Unbalanced Data

We developed AMV and ASV estimators of *p* and 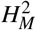 for analyses of balanced data because of their algebraic simplicity; however, phenotypic and genotypic data are nearly always unbalanced in practice. Here we extend the bias-correction solutions developed for balanced data to unbalanced data. ASV estimators of 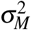 and 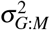 are developed here for a single marker locus (*M*) with unbalanced data. The phenotypic observations are entry-means 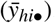.

The ASV estimator of the genetic variance associated with marker locus *M* for LMM (2) in its general form is:

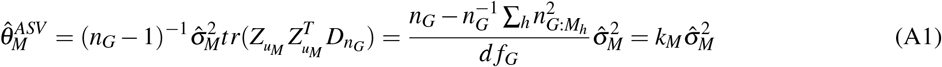

where 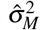 is the genetic variance associated with the marker locus effect, *d f*_*G*_ is the degrees of freedom for entries, and 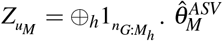 is a *k*_*M*_-bias corrected estimate of the marker-associated genetic variance. This form of *k*_*M*_ can be thought of as the general form of the correction term relative to (12) which is specific to balanced study designs.

The ASV estimate of the residual genetic variance among entries nested in the marker locus *M* is:

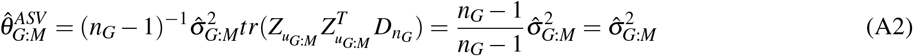

where 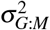 is the residual genetic variance among entries nested in the marker locus effect and 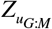 is a *n*_*G*_ identity matrix.

From equation (A1), the ASV estimator of *p* for a single marker locus (*M*) with unbalanced data is:

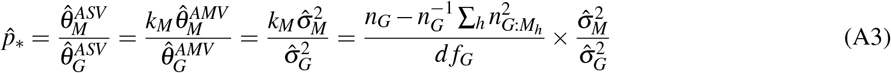

Hence, ASV yields unbiased estimates of the fraction of the genetic variance associated with the *M* locus (*p*) for unbalanced data. AMV estimates of *p* can be bias-corrected using the *k*_*M*_ coefficient shown above for unbalanced data. We review this here because most of the currently available linear mixed model software solutions produce AMV estimates, e.g., REML estimates of 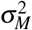 from LMM (2) are AMV estimates.

### A2 ASV Estimator of the Fraction of the Genetic Variance Associated with Two Marker Loci for Unbalanced Data

ASV estimators of 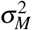 and 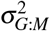 are developed here for two marker loci (*M*1 and *M*2). The phenotypic observations are entry-means 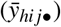, the underlying data are unbalanced, and the LMM for the entry-mean analysis is:

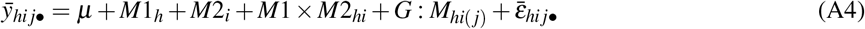

where 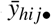 is the entry-mean, *µ* is the population mean, *h* = 1, 2, or 3, *i* = 1, 2, or 3, *j* = 1, 2, …, *n*_*G*_, *k* = 1, 2, …, *r*_*G*_, *M*1_*h*_ is the random effect of marker locus 1 with 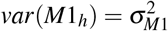, *M*2_*i*_ is the random effect of marker locus 2 with 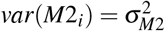, (*M*1 × *M*2)*hi* is the random effect of the interaction between marker loci 1 and 2 with 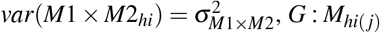 is the random effect of entries nested in marker loci with 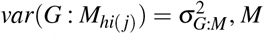, *M* refers to the *M*1 × *M*2 interaction, and 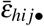 is the residual with 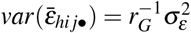.

The ASV estimator of the genetic variance associated with marker locus *M*1 from LMM (A4) is:

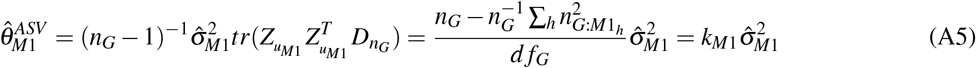

where 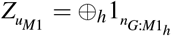 as is the incidence matrix for *M*1, 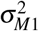 is the genetic variance associated with marker locus *M*1, *k*_*M*1_ is the bias-correction factor for *M*1. Similarly, the ASV estimator of the genetic variance associated with marker locus *M*2 from LMM (A4) is:

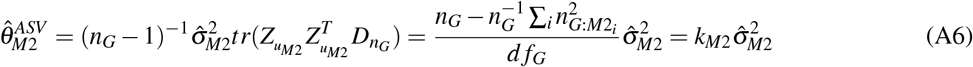

where 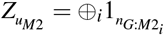 is the incidence matrix for *M*2.

The ASV estimator of the genetic variance associated with the interaction between marker loci *M*1 and *M*2 is:

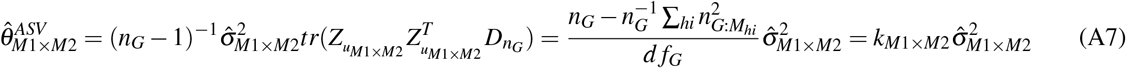

where 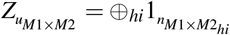 is the incidence matrix for the *M*1 by *M*2 interaction, 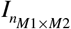 is a *n*_*M*1×*M*2_ identity matrix, 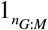 is a, *k*_*M*1×*M*2_ is the bias-correction factor for the interaction, 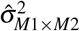 is the genetic variance associated with the interaction, and 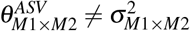.

The ASV estimator of the variance associated with the residual genetic variance among entries nested in marker loci *M*1 and *M*2 is:

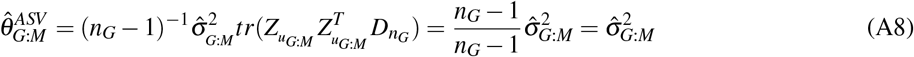

where 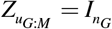 is a *n*_*G*_ incidence matrix.

Hence, as for the two-marker analysis, ASV yields *k*_*M*_-bias corrected estimates of the marker-associated genetic variance for two loci (*M*1 and *M*2) for unbalanced data:

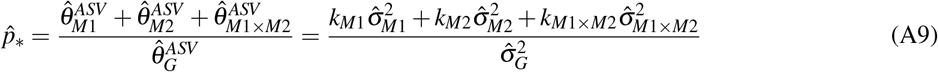

The genetic variance associated with marker loci can be partitioned into intra-locus (additive and dominance) and inter-locus (additive × additive, additive × dominance, and dominance × dominance) components [24, 25]. The example shown here includes all components: two degrees of freedom each for *M*1 and *M*2 and four degrees of freedom for *M*1 × *M*2.

From (A5) and (A6), the coefficients for bias-correcting AMV estimates of 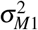 (the genetic variance explained by *M*1) and 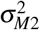 (the genetic variance explained by *M*2) are:

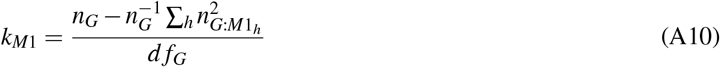

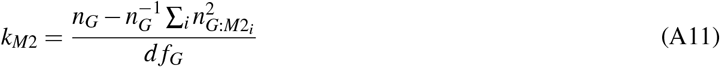

Similarly, from (A7), the coefficient for bias-correcting AMV estimates of 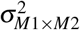 (the genetic variance explained by the interaction between *M*1 and *M*2) is:

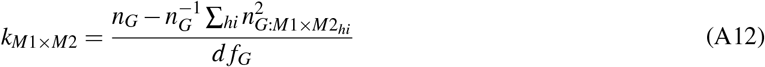

These *k*_*M*_-coefficients (A10-A12) can be substituted in (A9) to obtain bias-corrected estimates of *p*.

### A3 ASV Estimator of the Fraction of the Genetic Variance Associated with Three Marker Loci for Unbalanced Data

ASV estimators of the genetic variance associated with intra-locus and inter-locus effects of three marker loci are developed here for unbalanced data. As before, the phenotypic observations are entry-means 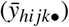 and the LMM for the entry-mean analysis is:

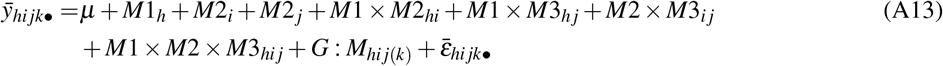

where 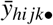 is the entry-mean, *µ* is the population mean, *h* = 1, 2, or 3, *i* = 1, 2, or 3, *j* = 1, 2, …, *n*_*G*_, *k* = 1, 2, …, *r*_*G*_, *M*1_*h*_ is the random effect of marker locus 1 with 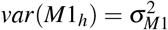, *M*2_*i*_ is the random effect of marker locus 2 with 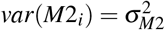*M*3_*j*_ is the random effect of marker locus 2 with 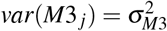, *M*1 × *M*2_*hi*_ is the random effect of the interaction between marker loci 1 and 2 with 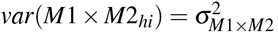, *M*1 × *M*3_*h j*_ is the random effect of the interaction between marker loci 1 and 3 with 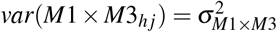, *M*2 × *M*3_*ij*_ is the random effect of the interaction between marker loci 2 and 3 with 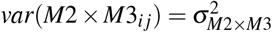 *M*1 × *M*2 × *M*3_*hi j*_ is the random effect of the interaction between marker loci 1, 2, and 3 with 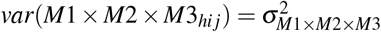, *G* : *M*_*hi j*(*k*)_ is the random effect of entries nested in marker loci with 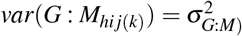, *M* refers to the *M*1 × *M*2 × *M*3 interaction, and 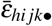 is the residual with 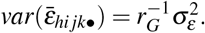.

The ASV estimator of the genetic variance associated with marker locus *M*1 from LMM (A13) is:

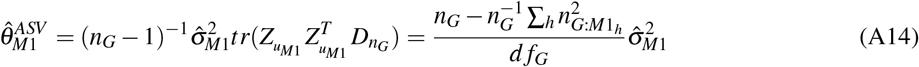

where 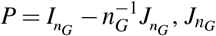 is a *n*_*G*_ unit matrix, 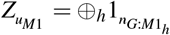 is the incidence matrix for *M*1. Substituting the appropriate values for marker loci *M*2 or *M*3 into (A14) yields the ASV estimators for these loci: 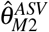 and 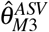.

The ASV estimator of the genetic variance associated with the two-locus interaction between marker loci *M*1 and *M*2 is:

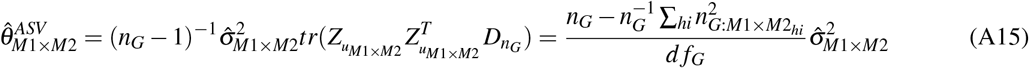

where 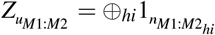 is the incidence matrix for *M*1 × *M*2 interaction. ASV estimates of the other two-locus interactions are similarly defined. The ASV estimator of the genetic variance associated with the three-locus interaction (*M*1 × *M*2 × *M*3) is:

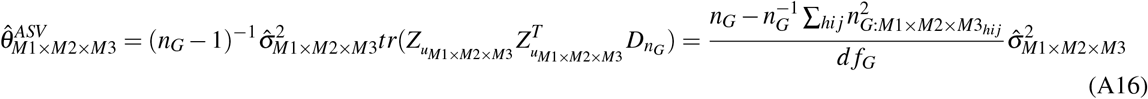

where 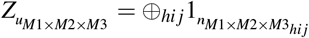 is the incidence matrix for the three-locus interaction. Finally, the ASV estimator of the residual genetic variance among entries nested in marker loci is:

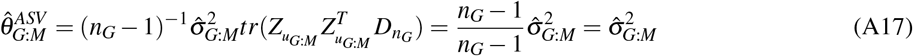

where 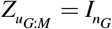 is a *n*_*G*_ identity matrix.

Hence, as for the three-marker analysis, ASV yields *k*_*M*_-bias corrected estimates of the marker-associated genetic variance for three loci (*M*1, *M*2, and *M*3) for unbalanced data:

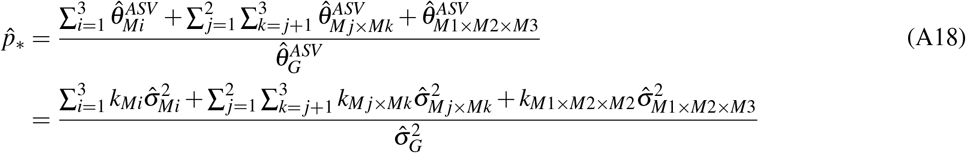

where *i* = 1, 2, 3, *j* = 1, 2, and *k* = 2, 3 and *j* ≠ *k* are used to specify *M*1, *M*2, *M*3 and all two-way locus-locus interactions.

From (A14), the *k*_*M*_-coefficient for bias-correcting AMV estimates of 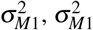, and 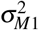 are:

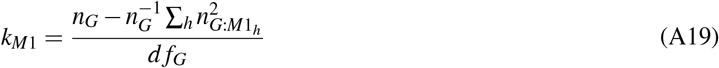

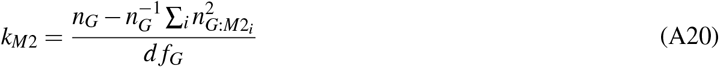

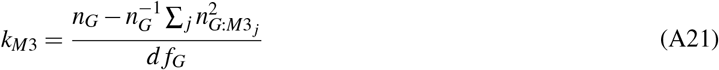

Similarly, from (A15), the *k*_*M*_-coefficient for bias-correcting AMV estimates of 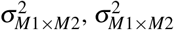, and 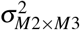 are:

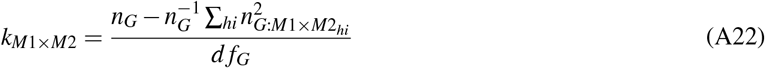

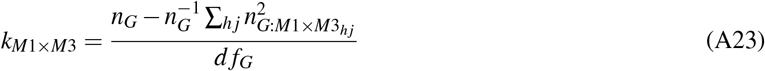

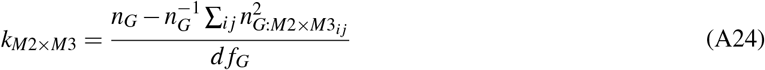

From (A16), the *k*_*M*_-coefficient for bias-correcting AMV estimates of 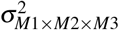 (the genetic variance explained by the three-way marker loci interaction) is:

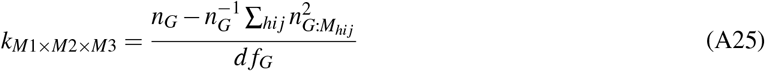

These *k*_*M*_-coefficients (A19-A25) are substituted in (A18) to obtain bias-corrected REML estimates of *p*.

### A4 Biases of AMV and ASV Estimators of Marker-Associated Variance

Following [68] and [125], the true value of the variance for the between-entry effect in LMM (1) is:

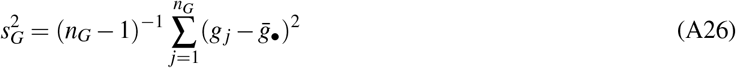

where 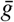 are the entry means, 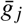 is the *j*^*th*^ element of 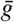, *j* = 1, 2, 3, …, *n*_*G*_, and 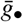 is the mean of 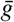. Note that 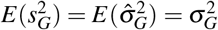. The variance of the residual effect in LMM (1) and (2) is:

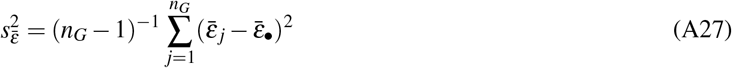

where 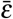 are the mean residuals across entries, 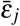 is the *j*^*th*^ element of 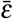, *j* = 1, 2, 3, …, *n*_*G*_, and 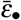 is the mean of 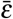. We note that 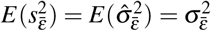. The variance for the entries nested in *M* effect in LMM (2) is:

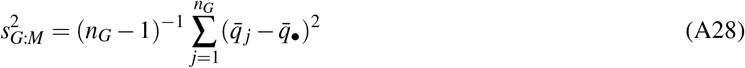

where 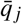 is the effect of the *j*^*th*^ entry nested in marker locus 1, *j* = 1, 2, 3, …, *n*_*G*_, and 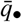 is the mean of 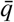 across entries. Note that 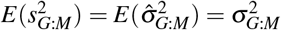. Finally, the variance for the effect of a single marker locus (*M*) in LMM (2) is:

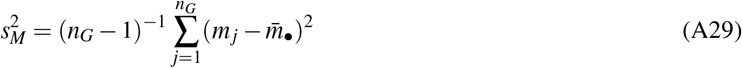

where *m*_*j*_ is the effect of the *j*^*th*^ genotype for a single marker locus, *j* = 1, 2, 3, …, *n*_*G*_, and 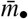 is the mean of *m* across entries. Note that the summation in (A29) is over entries and not marker alleles. We demonstrate in the main body of this text that 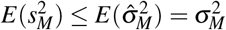 which causes *p* and 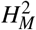 to be systematically overestimated. The expected value of 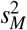 is:

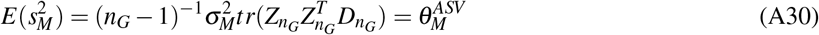

Note that the ASV estimator 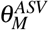 defines the expected value of 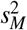 which, as we demonstrate in the main text, is a fraction of the AMV estimator 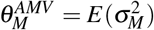. Our bias argument, supported by overwhelming simulation evidence, applies both in relation to the samples 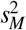 and 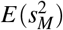, but we focus on the former as it is more tangible.

The bias of the AMV estimator of the marker-associated genetic variance is:

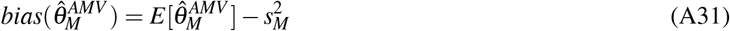

where 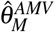is the AMV estimate of the marker associated genetic variance and 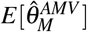 is the expected value of 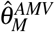. Similarly, the bias of the ASV estimator is:

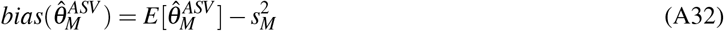

where 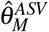 is the ASV estimator of the marker associated genetic variance and 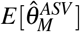 is the expected value of 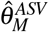.

The relative bias of the AMV estimator is:

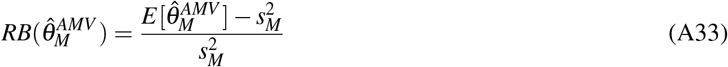

and the relative bias of the ASV estimator is:

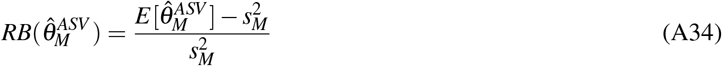

The biases and relative biases were empirically estimated through computer simulations.

### A5 Sample Variances for AMV and ASV Estimators of *p* and 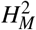

Sample variances for AMV and ASV estimators of *p* and 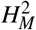 were developed using the delta method and basic covariance algebra [25, 122, 126]. The sample variance of the AMV estimator of *p* is:

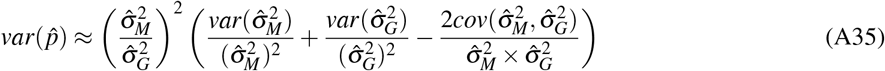

where 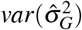 is the variance of 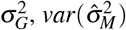 is the variance of 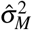, and 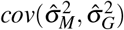 is the covariance between 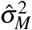 and 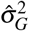. Similarly, the sample variance of the AMV estimator of 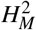 is:

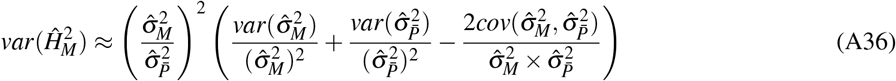

where 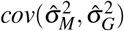 is the covariance between 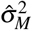 and 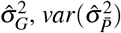 is the variance for the phenotypic variance on an entry-mean basis 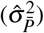, and 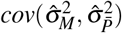 is the covariance between 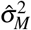 and 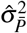 from equation (7). The variance of 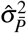 is:

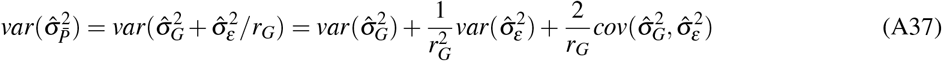

where 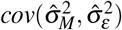 is the covariance between 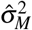 and 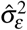. The covariance between 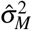 and 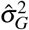 is:

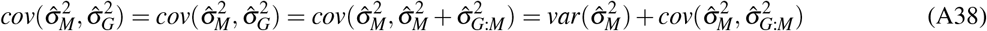

where 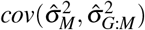 is the covariance between 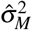 and 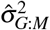. The covariance between 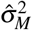 and 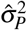 from equation (7) is:

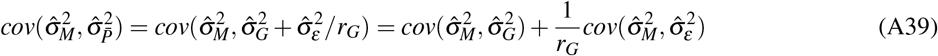

These sample variances can be extracted from the asymptotic variance-covariance matrices estimated with widely used software for linear mixed model analyses, e.g., *lme4* [78].

Using the delta method [25, 126] and recalling from equation (8) that 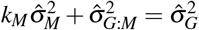, the sample variance of the ASV estimates of *p* is:

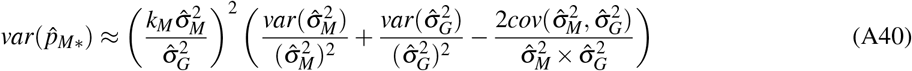

where 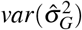 is the variance of 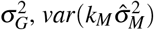 is the variance of 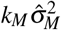, and 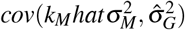 is the covariance between 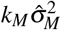 and 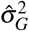.

Similarly, the sample variance for the ASV estimator of 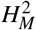 is:

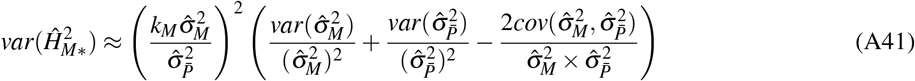

where 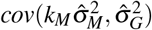 is the covariance between 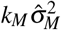 and 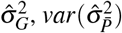 is the variance for the phenotypic variance on an entry-mean basis, and 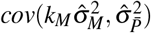 is the covariance between 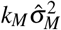 and 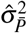 from equation (7). The covariance between 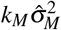 and 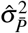 from equation (2) is:

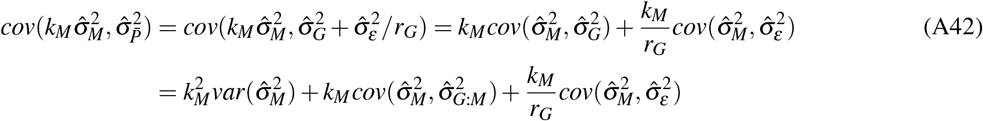

where 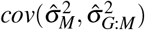 is the covariance between 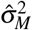 and 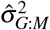 from equation (2). The covariance between 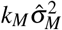 and 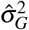 is:

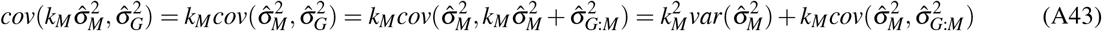

We found that the sample variances of the ASV estimators are consistently smaller than AMV estimators of *p* and 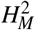 by a factor of 1 − *k*_*M*_, as shown here:

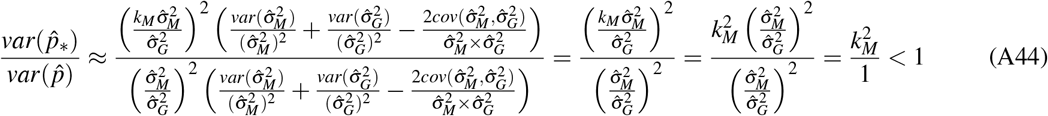

and

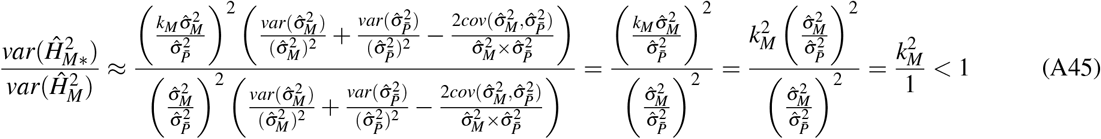

Importantly, 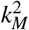 is the square of a value less than 1 indicating that the sampling variance of ASV estimates of *p*_∗_ and 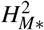 are less than those of *p* and 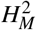.

**Table S1.**
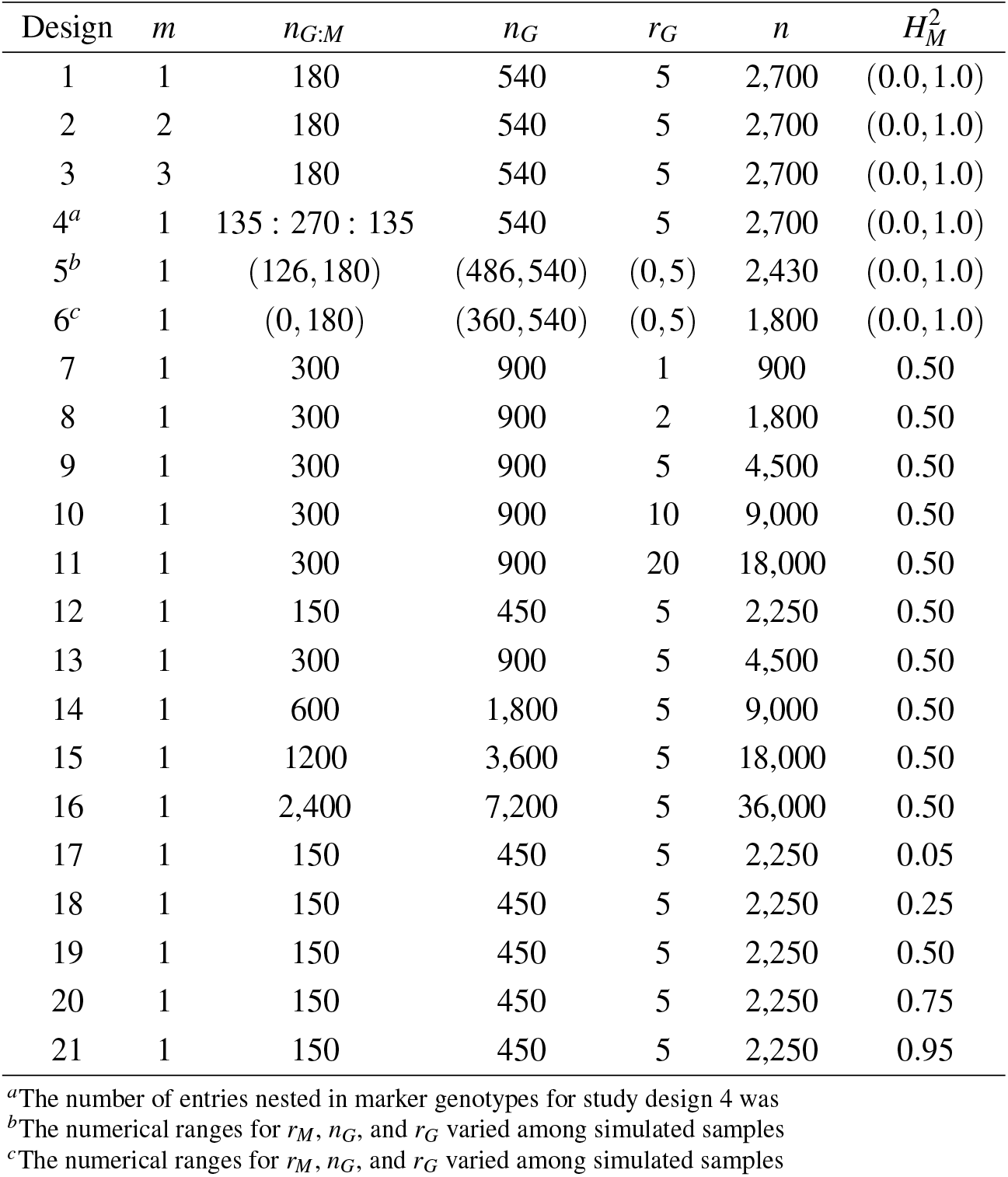
Simulation study designs and variables. Normally distributed phenotypic observations were simulated for 21 study designs and associated linear mixed models by varying the number of observations (*n* = *n*_*G*_ × *r*_*G*_), the number of entries (*n*_*G*_), the number of replications/entry (*r*_*G*_), the number of marker loci (*m*), *n*_*M*_ = 3 genotypes/marker locus, the number of entries/marker genotype (*n*_*G*:*M*_), and marker heritability 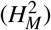. One thousand samples of size *n* were simulated for each study design. The segregation of a single marker locus in an F_2_ population was simulated in study design 4 The number of entries nested in marker genotypes for study design 4 was equivalent to the expected number for the segregation of a co-dominant DNA marker in a population segregating 1 *AA* : 2 *Aa* : 1 aa for a single marker locus. In this example, there are 135 entries nested in *AA*, 270 entries nested in *Aa*, and 135 entries nested in *aa* and each are replicated 5 times.simulates the segregation of a single locus in an F_2_ population The number of entries/genotype for study design 4

**Fig S1.**
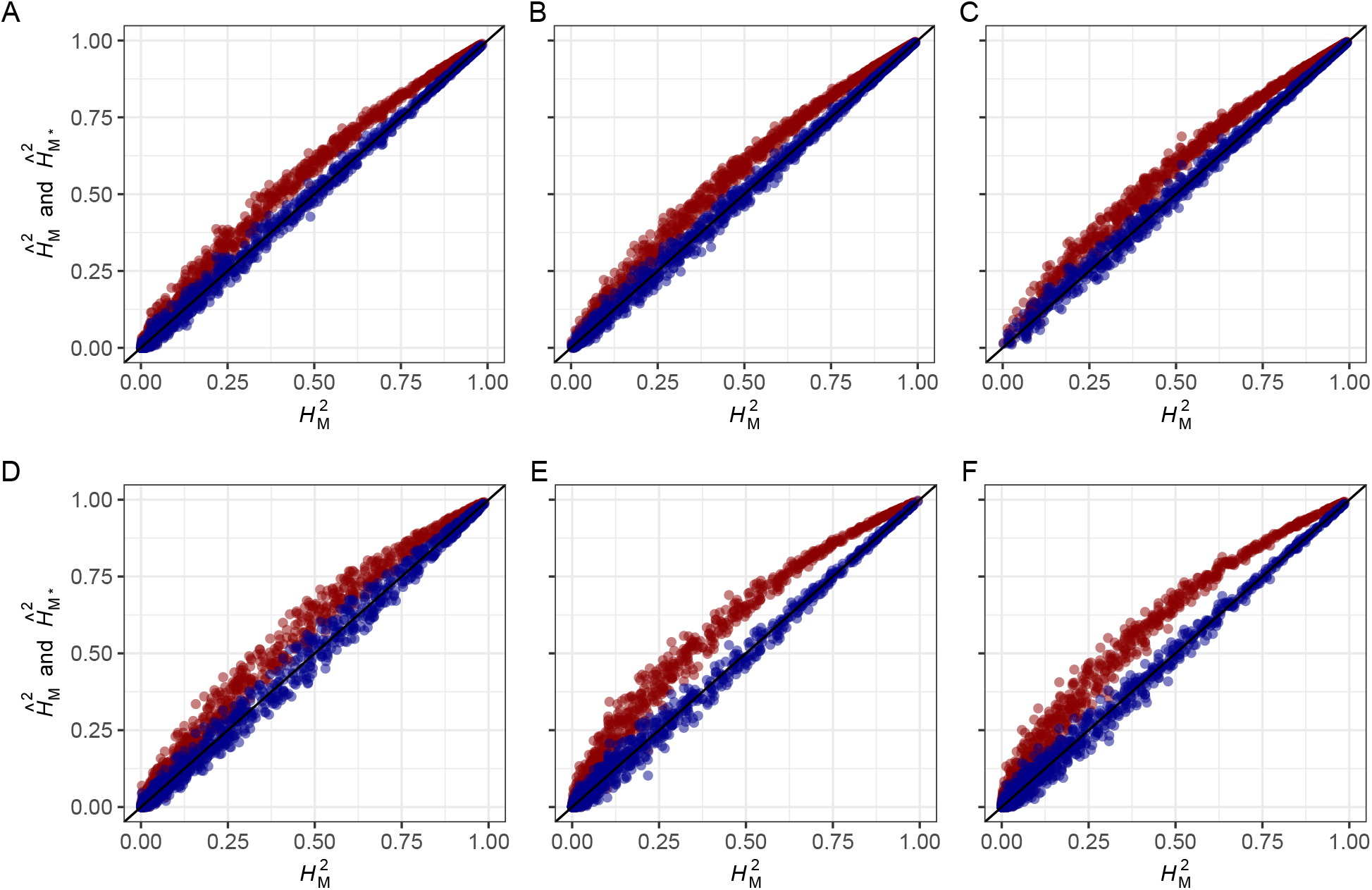
Accuracy of AMV and ASV Estimators of Marker Heritability When the Phenotypic Variance is Estimated by Pooling Marker and Residual Genetic Sources of Variation 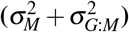. AMV and ASV estimates of 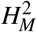 when 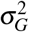 from LMM (1) is replaced with 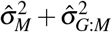 for AMV from LMM (2) or 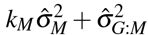 for ASV. Estimates are shown for 1,000 segregating populations simulated for different numbers of entries (*n*_*G*_ individuals, families, or strains), five replications/entry (*r*_*G*_ = 5), true marker heritability 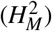 ranging from 0 to 1, and one to three marker loci with three genotypes/marker locus (*n*_*M*1_ = 3). The AMV estimates (shown in red) equal 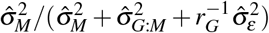, whereas the ASV estimates (shown in blue) equal 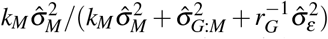. AMV estimates of marker heritability (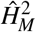; red highlighted observations) and ASV estimates of marker heritability (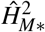; blue highlighted observations) are shown for: (A) one locus with balanced data for *n*_*G*_ = 540 entries (study design 1); (B) two marker loci with interaction (*M*1, *M*2, and *M*1 × *M*2) and balanced data for *n*_*G*_ = 540 (study design 2); (C) three marker loci with interactions (*M*1, *M*2, *M*3, *M*1 × *M*2, *M*1 × *M*3, *M*2 × *M*3, and *M*1 × *M*2 × *M*3) and balanced data for *n*_*G*_ = 540 (study design 3); (D) a population segregating 1:2:1 for a single marker locus with *r*_*G*:*M*_ = 135 entries for both homozygotes and *r*_*G*:*M*_ = 270 heterozygous entries, and *n*_*G*_ = 540 (study design 4); (E) one locus with 10% randomly missing data among 540 entries (study design 5); and (F) one locus with 33% randomly missing data among 540 entries (study design 6). Study design details are shown in Table S1.

**Fig S2.**
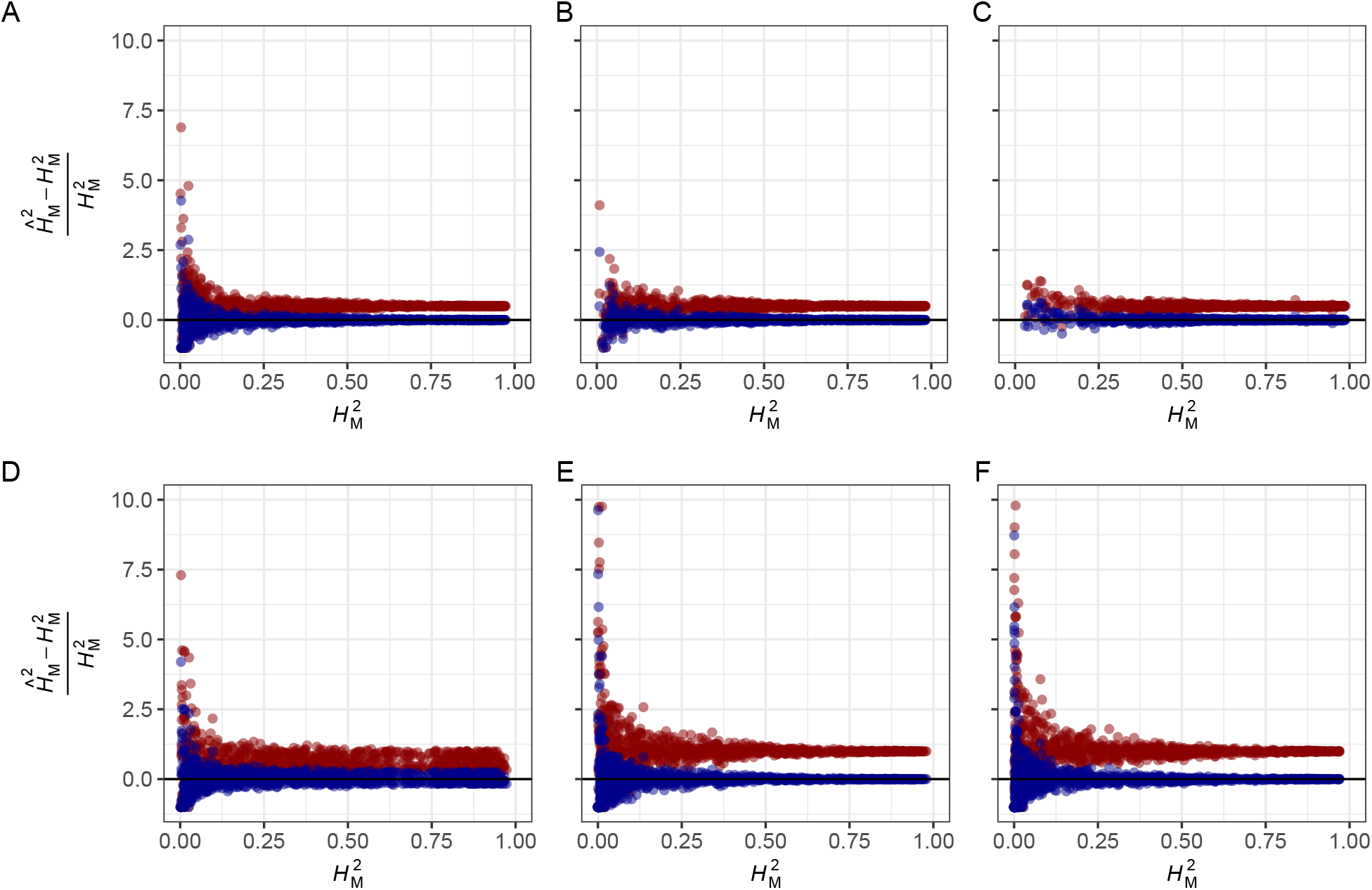
Relative Bias of AMV and ASV Estimators of Marker Heritability. Relative biases of AMV and ASV estimates of 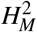 are shown for 1,000 segregating populations simulated for different numbers of entries (*n*_*G*_ individuals, families, or strains), five replications/entry (*r*_*G*_ = 5), true marker heritability 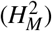 ranging from 0 to 1, and one to three marker loci with three genotypes/marker locus (*n*_*M*1_ = 3). AMV estimates of marker heritability (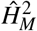; red highlighted observations) and ASV estimates of marker heritability (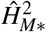 ; blue highlighted observations) are shown for: (A) one locus with balanced data for *n*_*G*_ = 540 entries (study design 1); (B) two marker loci with interaction (*M*1, *M*2, and *M*1 × *M*2) and balanced data for *n*_*G*_ = 540 (study design 2); (C) three marker loci with interactions (*M*1, *M*2, *M*3, *M*1× *M*2, *M*1 × *M*3, *M*2× *M*3, and *M*1 × *M*2× *M*3) and balanced data for *n*_*G*_ = 540 (study design 3); (D) an population segregating 1:2:1 for one marker locus with *r*_*G*:*M*_ = 135 entries for both homozygotes and *r*_*G*:*M*_ = 270 heterozygous entries, and *n*_*G*_ = 540 (study design 4); (E) one locus with 10% randomly missing data among 540 entries (study design 5); and (F) one locus with 33% randomly missing data among 540 entries (study design 6). Study design details are shown in Table S1.

